# Orthogonal RNA-regulated destabilization domains for three-color RNA imaging with minimal RNA perturbation

**DOI:** 10.1101/2024.12.16.628815

**Authors:** Tien G. Pham, Omoyemi Ajayi, Jiahui Wu

## Abstract

RNA-regulated destabilization domain allows live-cell imaging of single RNA. However, this method is limited to single-color RNA imaging and its long RNA tag induces destabilization of the tagged RNA. Here, we describe two orthogonal RNA-regulated destabilization domains that enable three-color mRNA imaging in living mammalian cells. One of these destabilizations can be combined with a short RNA tag (9XMS2) for single-molecule RNA imaging without perturbation of RNA stability.

## MAIN

RNA is one of the most fundamental biomolecules for gene expression regulation. Small nuclear RNAs can form spliceosome to generate mature mRNAs, mRNAs carry genetic information for protein synthesis, and microRNAs can fine-tune protein expression levels^1,2^. Therefore, robust methods for illuminating the spatiotemporal dynamics of different RNAs and their interactions in living cells are critical for understanding their roles in cellular function.

Conventional RNA imaging methods based on RNA hairpin tethering have high background fluorescence. In these methods, 24 repeats of an RNA hairpin, such as MS2, are fused to the 3’ untranslated region (3’UTR) of an RNA of interest^3^. Each of the MS2 RNA hairpins can bind and recruit fluorescent proteins (FPs) to the RNA of interest for imaging by MS2-MCP (MS2 coat protein) interactions^3^. However, the excess unbound FP-MCP create background fluorescence. To decrease background fluorescence, FP-MCP often needs to be knocked-in to host cells to ensure low expression levels. Although orthogonal RNA hairpin tethering methods have been reported^4^, knocking-in multiple FPs into host cell genome with proper expression levels can be highly laborious.

An alternative method to image RNA relies on fluorogenic RNA aptamers^5–11^. These fluorogenic RNA aptamers can bind and turn on the fluorescence signals of otherwise non-fluorescent small-molecule dyes^5–11^. When fused to the 3’UTR of an RNA of interest, these fluorogenic RNA aptamers can confer fluorescence signals to this RNA for imaging. Despite recent developments, many of the dyes for fluorogenic RNA aptamers have limited cell permeability and suboptimal photostability^12^. This limits the application of fluorogenic RNA aptamers for imaging cellular RNA with single-molecule resolution.

Recently, another RNA imaging method was developed based on an RNA-regulated destabilization domain^13^. In this method, an FP is fused to an RNA-regulated destabilization domain, termed tDeg. Because tDeg contains a degron sequence, this FP-tDeg fusion protein can be degraded rapidly in cells. However, FP-tDeg will only be stabilized when tDeg specifically binds to the Pepper RNA. By fusing multiple Peppers to the 3’UTR of an RNA of interest, this method enables imaging cellular RNA with single-molecule resolution^13^. Unlikely conventional RNA hairpin tethering method, FP-tDeg, can be introduced to host cells by transient transfecting its encoding DNA plasmids, making this method more accessible.

Although the Pepper-tDeg method allows imaging cellular RNA with single-molecule resolution, it still has two major limitations: (1) It only allows imaging of one RNA at a time in living cells due to a lack of orthogonal systems; (2) It requires a long RNA tag (1812 nt), which could destabilize the stability of the tagged RNA through nonsense-mediated mRNA decay^13,14^. The second problem is also shared by the other RNA imaging tags, such as the 24XBroccoli tag^15^ and the 24XMS2 tag^14^. Although tethering the poly(A) binding protein, PABPC1, or the polypeptide chain release factor, eRF3, to the 3’UTR of the tagged RNA can alleviate this problem^14^, matching the stoichiometry between the tethered protein to the tagged RNA, while avoiding overexpression of the tethered protein, could be challenging.

To address these two major limitations, we describe the development of two orthogonal systems to Pepper-tDeg for live-cell RNA imaging with minimal perturbation to RNA stability. These systems are based on the most widely used RNA-RNA binding protein pairs, MS2-MCP^3^ and PP7-PCP^4^, respectively. We converted the single-chain tandem MCP (stdMCP) and single-chain tandem PCP (stdPCP) to two orthogonal RNA-regulated destabilization domains, termed “mDeg” and “pDeg,” respectively. We showed that MS2-mDeg and PP7-pDeg enable imaging of cellular RNA with single-molecule resolution. Importantly, MS2-mDeg can image single mRNA much more effectively than the previously reported Pepper-tDeg system, representing a major improvement for RNA imaging. We also showed that MS2-mDeg, PP7-pDeg, and Pepper-tDeg are fully orthogonal to each other and they enable three-color simultaneous RNA imaging in living cells. Lastly, we developed a short MS2 tag (9XMS2) that can be used for mRNA imaging with minimal perturbation of RNA stability while retaining single-RNA sensitivity.

## RESULTS

### Development of mDeg, an MS2-regulated destabilization domain

To develop an orthogonal RNA-destabilization domain pair to Pepper-tDeg, we sought to convert stdMCP to an MS2-regulated destabilization domain. To achieve this, we sought to fuse a previously described C-terminal degron (Arg-Arg-Arg-Gly)^16,17^ to stdMCP such that, in the absence of MS2, this degron can be recognized by the E3 ubiquitin ligase complex to induce protein degradation. However, MS2 binding would block this degron from being recognized by the E3 ubiquitin ligase complex and thus inhibiting protein degradation. Since the C-terminus of stdMCP is far away from the MS2 binding site, we reasoned that appending the Arg-Arg-Arg-Gly degron to the C-terminus of stdMCP would unlikely to produce an MS2-regulated destabilization domain.

To convert stdMCP to an MS2-regulated destabilization domain, we therefore sought to circularly permute stdMCP to create a new C-terminus close to the MS2 binding site for appending the Arg-Arg-Arg-Gly degron. In this way, MS2 binding would block the Arg-Arg-Arg-Gly degron from recruitment of the proteasomal machinery needed for proteolysis. After close examination of the MS2-MCP crystal structure^18^ (PDB: 2BU1), we chose Arg49 and Arg83 of stdMCP for circular permutation because (1) both Arg are the same as the first amino acid of the Arg-Arg-Arg-Gly degron and we only needed to append Arg-Arg-Gly to extend this Arg to the full degron; (2) Both Arg are close to the MS2 binding site so that MS2 binding may block the Arg-Arg-Arg-Gly degron from recruitment of the proteasomal machinery needed for proteolysis. We therefore termed these two circularly permuted stdMCP with appended Arg-Arg-Gly as stdMCP-CP49-RRRG and stdMCP-CP83-RRRG, respectively (Supplementary Fig. 1).

We first asked whether fusing stdMCP-CP49-RRRG and stdMCP-CP83-RRRG, respectively, to a protein confers protein instability. To test this, we fused stdMCP-CP49-RRRG and stdMCP-CP83-RRRG to the C-terminus of enhanced yellow fluorescent protein (EYFP), respectively, and expressed this fusion protein in HEK293T cells. We found that cells expressing EYFP-stdMCP-CP49-RRRG and stdMCP-CP83-RRRG showed yellow fluorescence that is 50% and 2% of cells expressing EYFP, respectively (Supplementary Fig. 2). Together, these results suggest that both stdMCP-CP49-RRRG and stdMCP-CP83-RRRG can confer protein instability to EYFP. However, compared to stdMCP-CP49-RRRG, stdMCP-CP83-RRRG reduces the stability of EYFP to a much higher extent.

With the much higher destabilization effect from stdMCP-CP83-RRRG, we then asked whether this destabilization effect can be inhibited by binding to MS2. To test this, we coexpressed EYFP-stdMCP-CP83-RRRG with the MS2 RNA and a control RNA, in HEK293T cells. The MS2 RNA and the control RNA were expressed as circular RNA using the Tornado expression system^19^. We observed that, for EYFP-stdMCP-CP83-RRRG, coexpression with circular MS2 only led to a very limited increase in yellow fluorescence compared to the circular control RNA (Supplementary Fig. 3). This suggests that binding to MS2 did not effectively inhibit the Arg-Arg-Arg-Gly degron from inducing protein degradation. Together, these results indicate that MS2 cannot regulate the stability of stdMCP-CP83-RRRG.

Since stdMCP-CP83-RRRG’s stability cannot be regulated by MS2, we sought to further engineer stdMCP-CP49-RRRG to increase its destabilization effects. Even though stdMCP-CP49-RRRG has the same Arg-Arg-Arg-Gly degron as tDeg, stdMCP-CP49-RRRG only confers limited instability to EYFP (Supplementary Fig. 2). We reasoned that this decreased destabilization effects in stdMCP-CP49-RRRG might be due to the decreased accessibility of the Arg-Arg-Arg-Gly degron by the proteasomal machinery. Specifically, the Arg-Arg-Arg-Gly degron is fused to the flexible Tat peptide in tDeg, whereas the same degron is fused to a folded β-sheet in stdMCP-CP49-RRRG (Supplementary Fig. 1). This folded β-sheet may sterically limit the Arg-Arg-Arg-Gly degron from interacting with the proteasomal machinery.

We therefore asked whether adding a flexible linker between stdMCP-CP49 and the Arg-Arg-Arg-Gly degron can increase the accessibility of the degron by the proteasomal machinery and thus resulting in an increase in destabilization effect. To test this, we constructed DNA plasmids encoding EYFP-stdMCP-CP49-RRRG variants containing a flexible linker from one amino acid to six amino acids, respectively, between stdMCP-CP49 and the Arg-Arg-Arg-Gly degron (Fig. 1). Each EYFP-stdMCP-CP49-RRRG variant (termed mDeg variant) was expressed in HEK293T cells without the MS2 RNA. Compared to stdMCP-CP49-RRRG (mDeg variant0), we observed a gradual yellow fluorescence decrease from 28% to 92% as the linker length increases from mDeg variant1 to variant6, respectively (Fig. 1). Together, these results indicate that adding a flexible linker between stdMCP-CP49 and the Arg-Arg-Arg-Gly degron increases the degradation efficiency of the of the resulting destabilization domains.

**Fig. 1.**
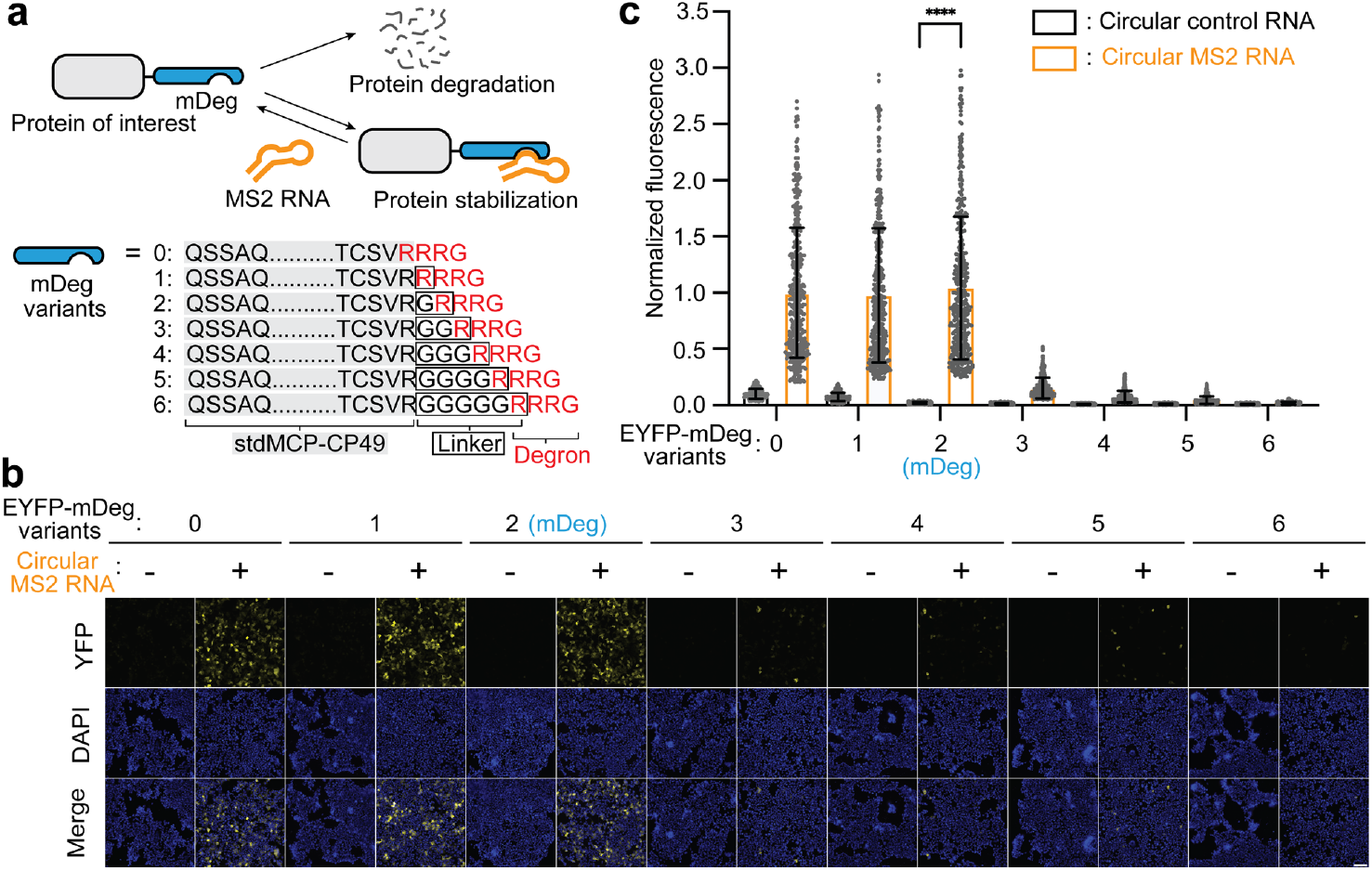
Development of mDeg, an MS2 RNA-regulated protein destabilization domain. **(a)** Schematic drawing of an MS2 RNA-regulated protein destabilization domain, mDeg. When fused to a protein of interest, mDeg (in blue) causes this fusion protein to be degraded in mammalian cells due to its C-terminal Arg-Arg-Arg-Gly degron. However, the protein destabilization function of mDeg is impeded when it binds to the MS2 RNA (in orange). To develop mDeg, we constructed seven mDeg variants (0-6) consisting of stdMCP-CP49 (highlighted in light grey) fused to a flexible linker with different lengths (highlighted in a black box) and the Arg-Arg-Arg-Gly degron (in red). **(b)** MS2 RNA stabilizes EYFP fused to mDeg (variant2) in HEK293T cells. To test whether MS2 can regulate the stability of different mDeg variants (0-6) shown in (**a**), we coexpressed each EYFP-mDeg variant with and without circular MS2 in HEK293T cells, respectively. While all the EYFP-mDeg variants can be regulated by MS2, we found that mDeg variant2 containing an “Arg-Gly” linker showed the highest yellow fluorescence increase of 55-fold upon binding to circular MS2. We termed variant2 as mDeg. All cells were stained with Hoechst’s dye. Scale bar, 100 μm. DAPI, 4′,6-diamidino-2-phenylindole. **(c)** Summary data of normalized fluorescence of each EYFP-mDeg variant with and without circular MS2 as in (**b**). Normalized average cellular yellow fluorescence of individual cells is plotted. (*n* = 3 independent cell cultures). Values are means ± s.d.

Next, we asked whether any of the above destabilization domains can be regulated by MS2. To test this, we coexpressed each EYFP-mDeg variant with and without circular MS2 in HEK293T cells, respectively. We found that all EYFP-mDeg variants can be regulated by MS2 but to different extents (Fig. 1). Among these, mDeg variant2 containing an “Arg-Gly” linker showed the highest yellow fluorescence increase of 55-fold upon binding to circular MS2 (Fig. 1). Thus, we chose this variant for further characterization and termed this MS2-regulated destabilization domain, “mDeg.”

### Development of pDeg, a PP7-regulated destabilization domain

Since stdMCP and stdPCP share high structural homology^4^, we therefore asked whether the design of mDeg can be applied to stdPCP for making a PP7-regulated destabilization domain. To test this, we constructed DNA plasmids encoding circularly permuted stdPCP variants at position 45 (equivalent to position 49 in stdMCP) with an appended linker followed by the Arg-Arg-Arg-Gly degron to the C-terminus (Supplementary Fig. 4). Similar to the strategy for designing mDeg, we titrated linker length from one to six amino acids between stdPCP-CP45 and the Arg-Arg-Arg-Gly degron (Supplementary Fig. 5). Each of these variants (termed pDeg variants) was coexpressed with and without circular PP7 in HEK293T cells. Similar to stdMCP-CP49-RRRG, stdPCP-CP45-RRRG (pDeg variant0) showed limited degradation efficiency as it exhibited 76% of yellow fluorescence compared to EYFP (Supplementary Fig. 5). The addition of linkers between stdPCP-CP45 and the Arg-Arg-Arg-Gly degron led to decreases of yellow fluorescence in the absence of circular PP7 (Supplementary Fig. 5). When PP7 was coexpressed, we found that pDeg variant2 containing an “Arg-Gly” linker showed the highest fold change in yellow fluorescence of 22-fold (Supplementary Fig. 5). Unexpectedly, we also observed yellow-fluorescent puncta in all pDeg variants (Supplementary Fig. 6). Together, these results suggest that pDeg variant2 can serve as a destabilization domain and its destabilization effect can be inhibited when bound by PP7.

Even though pDeg variant2 can be regulated by the PP7 RNA, the yellow-fluorescent puncta suggest aggregation of the EYFP-pDeg variant2-PP7 complex, which could generate artifacts when used for RNA imaging. We therefore sought to eliminate this aggregation phenomenon of the EYFP-pDeg variant2-PP7 complex.

We first asked whether the yellow-fluorescent puncta of EYFP-pDeg variant2 is due to aggregation of the circular PP7 RNA. To test this, we transiently expressed EYFP-pDeg variant2 lacking the Arg-Arg-Arg-Gly degron (termed pDeg variant7) in HEK293T in the absence of circular PP7 RNA. Without the Arg-Arg-Arg-Gly degron, EYFP-pDeg variant7 should be stable in the absence of the circular PP7 RNA. If the yellow-fluorescent puncta are due to the circular PP7 RNA, we should not observe any of these puncta in cells expressing EYFP-pDeg variant7. However, we found that EYFP-pDeg variant7 showed similar yellow-fluorescent puncta as in cells expressing the EYFP-pDeg variant2-PP7 complex (Supplementary Fig. 6 and 7). Thus, these results suggest that the aggregation of the EYFP-pDeg variant2-PP7 complex does not come from the circular PP7 RNA.

Circular permutation could change the energetics of transient partial unfolding in proteins, which may lead to protein aggregation^20^. We therefore asked whether the yellow-fluorescent puncta in cells expressing the EYFP-pDeg variant2-PP7 complex are due to aggregation of pDeg variant2. To test this, we constructed a DNA plasmid encoding a non-circularly permuted stdPCP fused to EYFP and expressed this in HEK293T cells. If circular permutation is the cause of the yellow-fluorescent puncta, such puncta will not be expected in cells expressing EYFP-stdPCP. Unexpectedly, we found that cells expressing EYFP-stdPCP showed the same yellow-fluorescent puncta as in cells expressing the EYFP-pDeg variant2-PP7 complex (Supplementary Fig. 6 and 7). Thus, these results suggest that the yellow-fluorescent puncta are not caused by circular permutation of stdPCP.

Wild-type PCP can oligomerize to form capsids for the *Pseudomonas* bacteriophage via interactions between a loop region, called the FG loop^21^. Although we used a capsid assembly-deficient PCP variant with part of its FG loop (Cys67 to Phe74) deleted when constructing the pDeg variants, we nevertheless asked whether the remaining FG loop residues could be the cause of aggregation. To test this, we replaced the remaining FG loop (Val64 to Val78) with a single Gly residue in pDeg variant2 and termed this pDeg variant2(del loop). We then coexpressed EYFP-pDeg variant2(del loop) with and without circular PP7 in HEK293T cells.

Without PP7, only minimal yellow fluorescence above background fluorescence was detected. However, when PP7 was coexpressed, cells exhibited a 21-fold increase of yellow fluorescence (Fig. 2). Furthermore, we did not observe any yellow-fluorescent puncta in the EYFP-pDeg variant2(del loop)-PP7 complex (Supplementary Fig. 8). Together, these results suggest that the fluorescent puncta of EYFP-pDeg variant2 was caused by the remaining residues in the FG loop. Since the stability of pDeg variant2 (del loop) can be regulated by PP7, we therefore termed this PP7-regulated destabilization domain, “pDeg.”

**Fig. 2.**
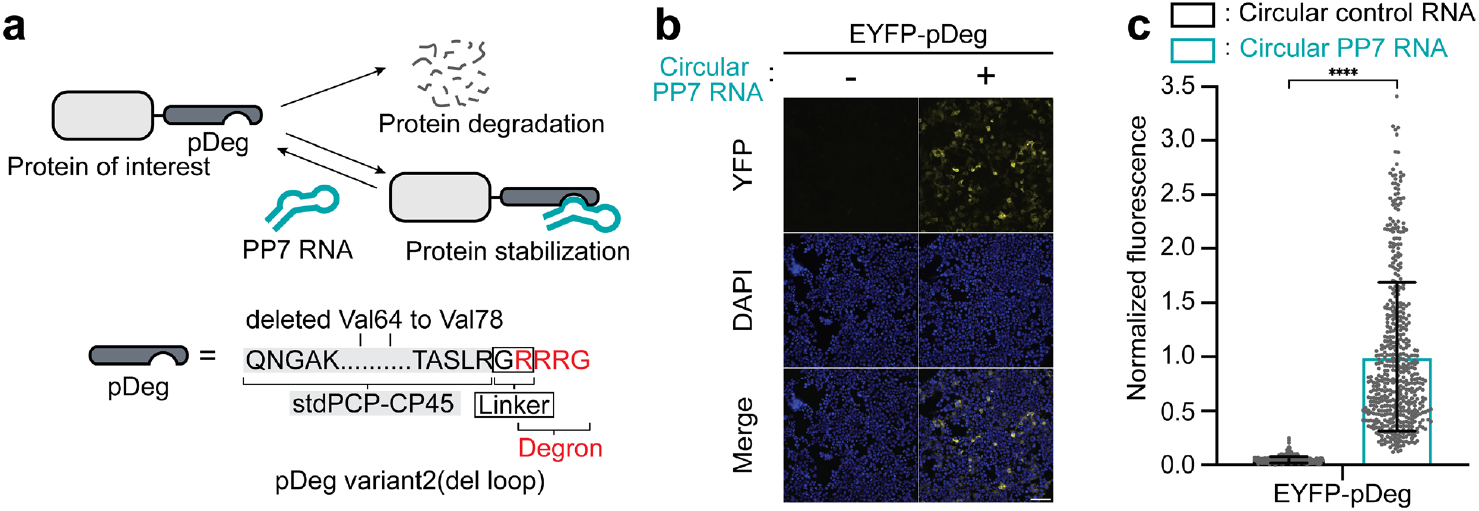
Development of pDeg, a PP7 RNA-regulated protein destabilization domain. **(a)** Schematic drawing of a PP7 RNA-regulated protein destabilization domain, pDeg. Similar to mDeg, pDeg (in dark grey) induces protein degradation when fused to a protein of interest. However, binding to the PP7 RNA (in tiffany green) inhibits the protein destabilization function of pDeg. pDeg consists of stdPCP-CP45 with the FG loops (Val64 to Val78) deleted (highlighted in light grey) fused to a ‘Gly-Arg’ linker (highlighted in a black box) and the Arg-Arg-Arg-Gly degron (in red). **(b)** PP7 RNA stabilizes EYFP fused to pDeg in HEK293T cells. To test whether PP7 can regulate the stability of pDeg, we coexpressed EYFP-pDeg with and without circular PP7 in HEK293T cells. Without PP7, only minimal yellow fluorescence above background fluorescence was detected. However, when circular PP7 was coexpressed, cells exhibited a 21-fold increase of yellow fluorescence. All cells were stained with Hoechst’s dye. Scale bar, 100 μm. **(c)** Summary data of normalized fluorescence of EYFP-pDeg with and without circular PP7 as in (**b**). Normalized average cellular yellow fluorescence of individual cells is plotted. (*n* = 3 independent cell cultures). Values are means ± s.d.

### MS2-mDeg and PP7-pDeg can regulate the stability of various fluorescent proteins

Next, we asked whether MS2-mDeg and PP7-pDeg can regulate other fluorescent proteins besides EYFP. To test this, we fused mDeg to the C-terminus of tdStayGold^22^, mCherry^23^, and HaloTag^24^, and pDeg to the C-terminus of mEGFP^25^, mScarlet-I3^26^, and iRFP670^27^, respectively. Each of these proteins was transiently expressed in HEK293T cells with and without its cognate RNA (i.e. MS2 or PP7). Additionally, we incubated cells expressing HaloTag-mDeg with the JF646 ligand^28^ prior to imaging. The expression levels of each protein were measured by fluorescence. In each case, there was a considerable increase in fluorescence of the mDeg-tagged and pDeg-tagged protein only when their cognate RNA was coexpressed in cells (Fig. 3 and Supplementary Fig. 9). Together, these results showed that MS2-mDeg and PP7-pDeg can regulate a diverse array of fluorescent proteins and the HaloTag.

**Fig. 3.**
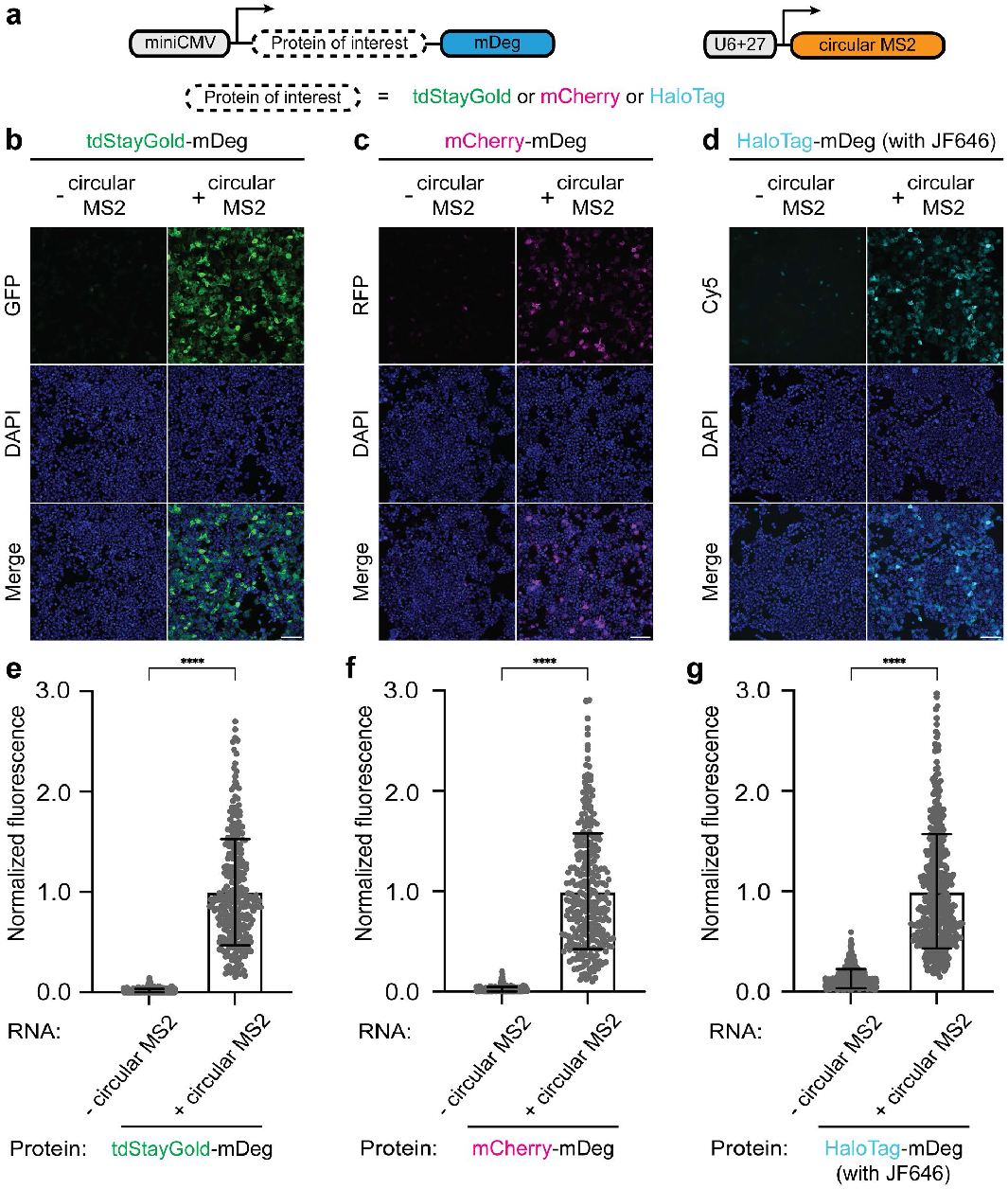
MS2-mDeg confers MS2-dependent regulation to various fluorescent proteins and HaloTag. (**a**) Scheme of plasmids that encode protein of interest fused to mDeg, and circular MS2 RNA, respectively. Protein of interest includes tdStayGold, mCherry, and HaloTag. (**b**-**d**) MS2-mDeg can regulate the stability of various fluorescent proteins and HaloTag. To test whether the MS2 RNA stabilizes different proteins fused to mDeg, we imaged HEK293T cells expressing tdStayGold (**b**), mCherry (**c**), and HaloTag (**d**) fused to mDeg with and without circular MS2, respectively. In each case, there was a considerable increase in fluorescence. For imaging HaloTag-mDeg, cells were incubated in media with the JF646 ligand. All cells were stained with Hoechst’s dye. Scale bar, 100 μm. (**e**-**g**) Summary data of normalized fluorescence of mDeg-fused fluorescent proteins and HaloTag with and without circular MS2 as in (**b**-**d**). Normalized average cellular fluorescence of individual cells is plotted. (*n* = 3 independent cell cultures). Values are means ± s.d.

### MS2-mDeg and PP7-pDeg can regulate protein stability in different cell types

We then asked if the stability of EYFP-mDeg and EYFP-pDeg can be regulated by MS2 and PP7, respectively in different mammalian cell lines. To test this, we expressed EYFP-mDeg and EYFP-pDeg in U2OS, COS-7, and HeLa cells, respectively. In each case, the EYFP-mDeg and EYFP-pDeg were coexpressed with and without their cognate RNA. Similar to what we observed in HEK293T cells, EYFP-mDeg and EYFP-pDeg showed minimal fluorescence in all three cell lines without its cognate RNA (Supplementary Fig. 10 and 11). However, the yellow fluorescence intensity of EYFP-mDeg and EYFP-pDeg was considerably increased only when the cognate RNA was coexpressed (Supplementary Fig. 10 and 11). Taken together, these data indicate that protein stability of EYFP-mDeg and EYFP-pDeg can be regulated by MS2 and PP7, respectively, in different mammalian cell lines.

### MS2-mDeg and PP7-pDeg can image single RNA in living cells

MS2-MCP has been one of the earliest methods for live-cell RNA imaging and has remained the gold standard over the past two decades^3,29,30^. However, one limitation of this system is the high levels of background fluorescence generated by the excess EGFP-MCP. To decrease background fluorescence, EGFP-MCP needs to be fused to a nuclear localization sequence (NLS) for sending the unbound EGFP-MCP from the cytosol to the nucleus^3^. However, it has been suggested that potential artifacts may occur since each MS2-tagged mRNA will contain dozens of NLS due to the recruited NLS-EGFP-MCP^31^. Additionally, to further decrease background fluorescence, NLS-EGFP-MCP often needs to be stably introduced to the host cell genome, which is a technical hurdle for using this system.

Since fluorogenic proteins containing mDeg and pDeg can be rapidly degraded in cells when not bound to their cognate RNA, this system provides a potential alternative for mRNA imaging without the need for NLS and stable cell lines. We therefore asked if mDeg-fused and pDeg-fused fluorogenic proteins can image mRNA in living cells by transient transfection. To test this, we transiently expressed (mNeonGreen)_4_-mDeg and (mNeonGreen)_4_-pDeg in U2OS cells to image an *ER-targeted mTagBFP2* reporter mRNA (*CyTERM-mTagBFP2*)^Ref,32^ containing a previously reported 24XMS2 tag and 24XPP7 tag in the 3’UTR^33,34^, respectively. In both cases, we observed green-fluorescent puncta, likely reflecting single *mTagBFP2* mRNA (Fig. 4). In addition, these green-fluorescent puncta were not observed in control cells expressing the same *mTagBFP2* reporter mRNA without any 24XMS2 tag or 24XPP7 tag (Fig. 4). This indicates that the green-fluorescent puncta were from (mNeonGreen)_4_-mDeg and (mNeonGreen)_4_-pDeg bound to the 24XMS2 tag and 24XPP7 tag, respectively. Taken together, these results suggest that (mNeonGreen)_4_-mDeg and (mNeonGreen)_4_-pDeg can track the dynamics of single mRNA in living cells by transient transfection.

**Fig. 4.**
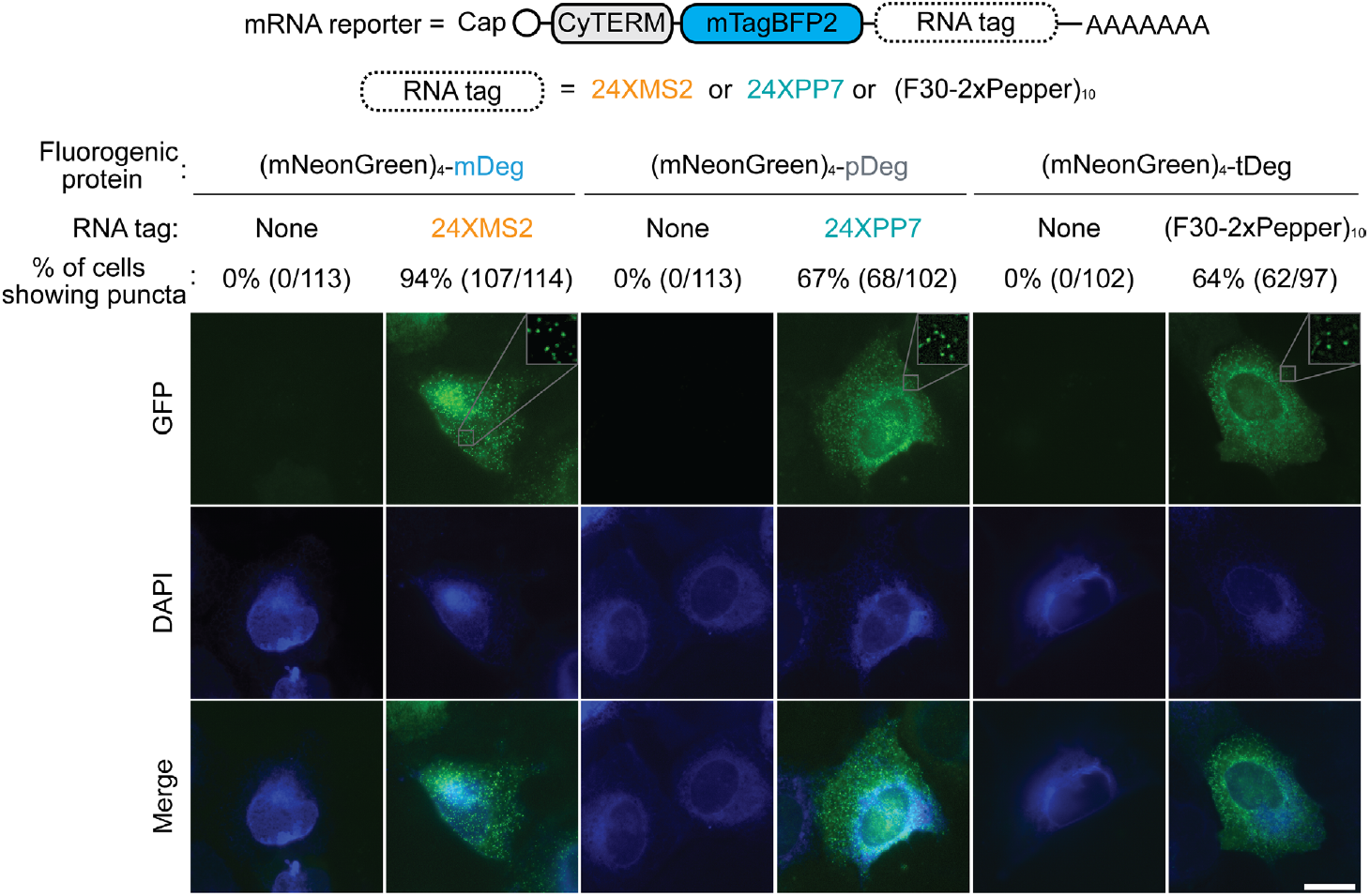
Comparison of MS2-mDeg and PP7-pDeg to existing fluorogenic system, Pepper-tDeg, for mRNA imaging. To compare the performance of MS2-mDeg, PP7-pDeg, and Pepper-tDeg for mRNA imaging, we imaged cells expressing an *mTagBFP2* reporter mRNA tagged with 24XMS2, 24XPP7, and (F30-2xPepper)_10_, respectively. We imaged these tagged *mTagBFP2* mRNA by coexpressing fluorogenic proteins consisting of (mNeonGreen)_4_ fused to the cognate RNA-regulated destabilization domain, i.e. mDeg, pDeg, and tDeg, respectively. In each case, we observed cytosolic green-fluorescent puncta, reflecting the *mTagBFP2* reporter mRNA. Among these three MS2-mDeg can detect cellular mRNA most effectively 94% (107 out of 114 transfected cells), while pDeg-PP7 (67%, 68 out of 102 cells) performed on par with the previously reported tDeg-Pepper system (64%, 62 out of 97 cells). Scale bar, 20 μm.

### Comparison of MS2-mDeg, PP7-pDeg, and Pepper-tDeg for mRNA imaging

We next asked whether mDeg-MS2 and pDeg-PP7 can detect mRNA in living cells as effectively as by the previously reported Pepper-tDeg method^13^. To test this, we generated DNA plasmids encoding the same ER-targeted mTagBFP2 reporter mRNA containing 24XMS2, 24XPP7, and (F30-2xPepper)_10_, respectively, in the 3’UTR. We imaged these tagged *mTagBFP2* mRNA by coexpressing (mNeonGreen)_4_ fused to the cognate RNA-regulated destabilization domain in U2OS cells. To assess the performance of each RNA imaging system, we quantified the percentage of transfected cells showing green-fluorescent puncta in the cytosol. Among these three systems, we observed the highest percentage of transfected cells showing green-fluorescent puncta using mDeg-MS2 94% (107/114 cells) (Fig. 4). Compared to mDeg-MS2, we observed a lower percentage of transfected cells showing green-fluorescent puncta with pDeg-PP7 (67%, 68/102 cells) and tDeg-Pepper (64%, 62/97 cells) (Fig. 4). Together, these results suggest that mDeg-MS2 can detect mRNA most effectively, while pDeg-PP7 performed on par with the previously reported tDeg-Pepper system.

### MS2-mDeg, PP7-pDeg, and Pepper-tDeg are orthogonal to each other

We next asked whether MS2-mDeg, PP7-pDeg, and Pepper-tDeg are orthogonal to each other since orthogonality between the three RNA-protein pairs is a prerequisite for using them to image multiple RNA simultaneously in the same cell. To test pairwise orthogonality, we coexpressed all nine possible combinations of EYFP-mDeg, EYFP-pDeg, and EYFP-tDeg with their cognate and non-cognate RNA in HEK293T cells, respectively (Fig. 5a, b). If MS2-mDeg, PP7-pDeg, and Pepper-tDeg are fully orthogonal, we would expect to observe yellow fluorescence only when an RNA-regulated destabilization domain is coexpressed with its cognate RNA. Indeed, we only observed strong yellow fluorescence when an EYFP-fused destabilization domain was coexpressed with its cognate RNA (Fig. 5a, b). Coexpression of each EYFP-fused destabilization domain with a non-cognate RNA showed minimal yellow fluorescence in cells (Fig. 5a, b). Together, these results suggest that MS2-mDeg, PP7-pDeg, and Pepper-tDeg are fully orthogonal in mammalian cells.

**Fig. 5.**
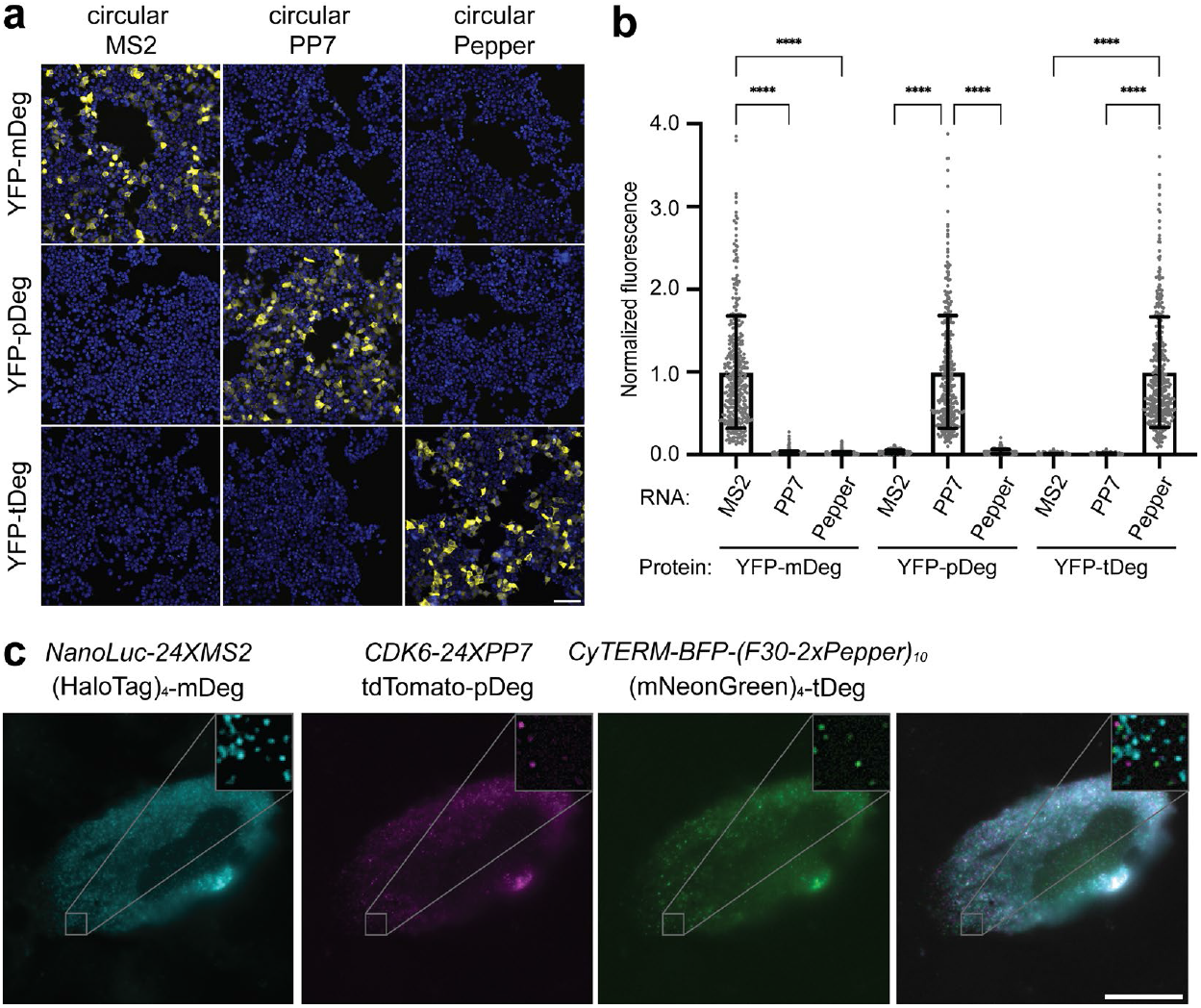
MS2-mDeg, PP7-pDeg, and Pepper-tDeg are orthogonal to each other and they enable three-color RNA imaging in living cells. **(a)** MS2-mDeg, PP7-pDeg, and Pepper-tDeg are orthogonal to each other. To test the orthogonality between MS2-mDeg, PP7-pDeg, and Pepper-tDeg, we coexpressed all nine possible combinations of EYFP-mDeg, EYFP-pDeg, and EYFP-tDeg with their cognate and non-cognate RNA in HEK293T cells, respectively. We only observed strong yellow fluorescence when an EYFP-fused destabilization domain was coexpressed with its cognate RNA, suggesting that these three systems are fully orthogonal in mammalian cells. All cells were stained with Hoechst’s dye. Scale bar, 100 μm. **(b)** Summary data of normalized fluorescence of (**a**) Average cellular fluorescence of EYFP-fused destabilization domain with non-cognate RNA is normalized to the same protein with cognate RNA. (*n* = 3 independent cell cultures). Values are means ± s.d. **(c)** MS2-mDeg, PP7-pDeg, and Pepper-tDeg enable three-color RNA imaging in living cells. To simultaneously image three different RNA species in U2OS cells, we coexpressed three mRNA reporters: *Nanoluc-24XMS2, CDK6-24XPP7*, and *CyTERM-mTagBFP2-(F30-2xPepper)*_*10*_ with three fluorogenic proteins: (HaloTag)_4_-mDeg (with the JF646 ligand), tdTomato-pDeg, and (mNeonGreen)_4_-tDeg. We observed mobile fluorescent puncta in the cytosol reflecting cellular mRNAs as shown in Supplementary Video1. Scale bar, 20 μm.

### MS2-mDeg, PP7-pDeg, and Pepper-tDeg enable three-color RNA imaging in living cells

We next sought to image multiple different RNAs in living cells at the same time using the orthogonal RNA-regulated destabilization domains. We first started by imaging two different RNAs in living mammalian cells. For two-color live-cell mRNA imaging, we expressed two reporter plasmids expressing a *Nanoluc* tagged with 24XMS2 and a *CyTERM-mTagBFP2* tagged with 24xPP7, respectively. In the same cells, we coexpressed (HaloTag)_4_-mDeg (with the JF646 ligand) and (AausGFP)_2_-pDeg, a GFP with high molecular brightness^35^, as fluorogenic proteins. We observed distinct but non-overlapping fluorescent puncta in both the GFP channel and Cy5 channel, respectively, reflecting individual mRNAs (Supplementary Fig. 12). These results suggest that MS2-mDeg and PP7-pDeg can be used for tracking the dynamics of two RNA species in the same cell at the same time.

After validating that MS2-mDeg and PP7-pDeg can be used for two-color RNA imaging, we then sought to simultaneously image three different RNA species in living cells by combining them with the Pepper-tDeg system. To test this, we coexpressed three reporter plasmids expressing a *Nanoluc* mRNA tagged with 24XMS2, a *CDK6* mRNA tagged with 24XPP7, and a *CyTERM-mTagBFP2* tagged with (F30-2xPepper)_10_ tag in U2OS cells. To visualize these reporter mRNAs, we coexpressed (HaloTag)_4_-mDeg (with the JF646 ligand), tdTomato-pDeg, and (mNeonGreen)_4_-tDeg as fluorogenic proteins. In this case, we again observed fluorescent puncta in the cytosol reflecting cellular mRNAs (Fig. 5c and Supplementary Video 1). Together, these results suggest that MS2-mDeg, PP7-pDeg, and Pepper-tDeg can be used for simultaneous imaging of three different mRNA species in living cells.

### MS2-mDeg can be combined with photostable fluorogenic proteins for continuous RNA imaging for an extended period

Since MS2-mDeg has the best performance for single RNA imaging among the three pairs, we asked whether mDeg can be combined with photostable fluorophores for continuous RNA imaging for an extended period of time. To test this, we chose (HaloTag)_4_-mDeg with the JF646 ligand as the fluorogenic protein due to its high photostability. We used this fluorogenic protein to image a *NanoLuc-24XMS2* mRNA in living U2OS cells with continuous excitation and image acquisition. We observed mobile far-red fluorescent puncta with continuous image acquisition for 120 seconds with limited photobleaching (Supplementary Video 2). These results indicate that MS2-mDeg can be combined with photostable fluorogenic proteins for live-cell RNA imaging with continuous image acquisition for an extended period of time.

### Development of short RNA imaging tags with minimal perturbation of RNA stability

A recent study has shown that the 24XMS2 tag can destabilize the tagged mRNA through nonsense-mediated mRNA decay due to its excess length^14^. Since MS2-mDeg is a fluorogenic system and can detect mRNA more effectively than PP7-pDeg and Pepper-tDeg (Fig. 4). Therefore, we reasoned that we might be able to use a shorter MS2 RNA tag to image mRNA without perturbing its stability.

To design a short MS2 tag with minimal perturbation to RNA stability, we first sought to identify the minimal repeat number of MS2 that does not destabilize the tagged mRNA. To test this, we constructed reporter plasmids expressing *mCherry* mRNA with different repeat numbers of MS2 at the 3’UTR, including 3X, 6X, 9X, 12X, and 24X respectively. We used RT-qPCR to quantify the levels of the reporter mRNA transcripts in HEK293T cells. Consistent with previous report^13,14^, 24XMS2 destabilized the tagged *mCherry* mRNA levels by over 50% compared to an untagged reporter mRNA (Fig. 6a). However, we also found that 9XMS2 has a minimal effect on the stability of the tagged *mCherry* mRNA (Fig. 6a). Together, these results suggest that a short imaging tag containing 9XMS2 has minimal perturbation on the tagged *mCherry* mRNA’s stability in living cells.

**Fig. 6.**
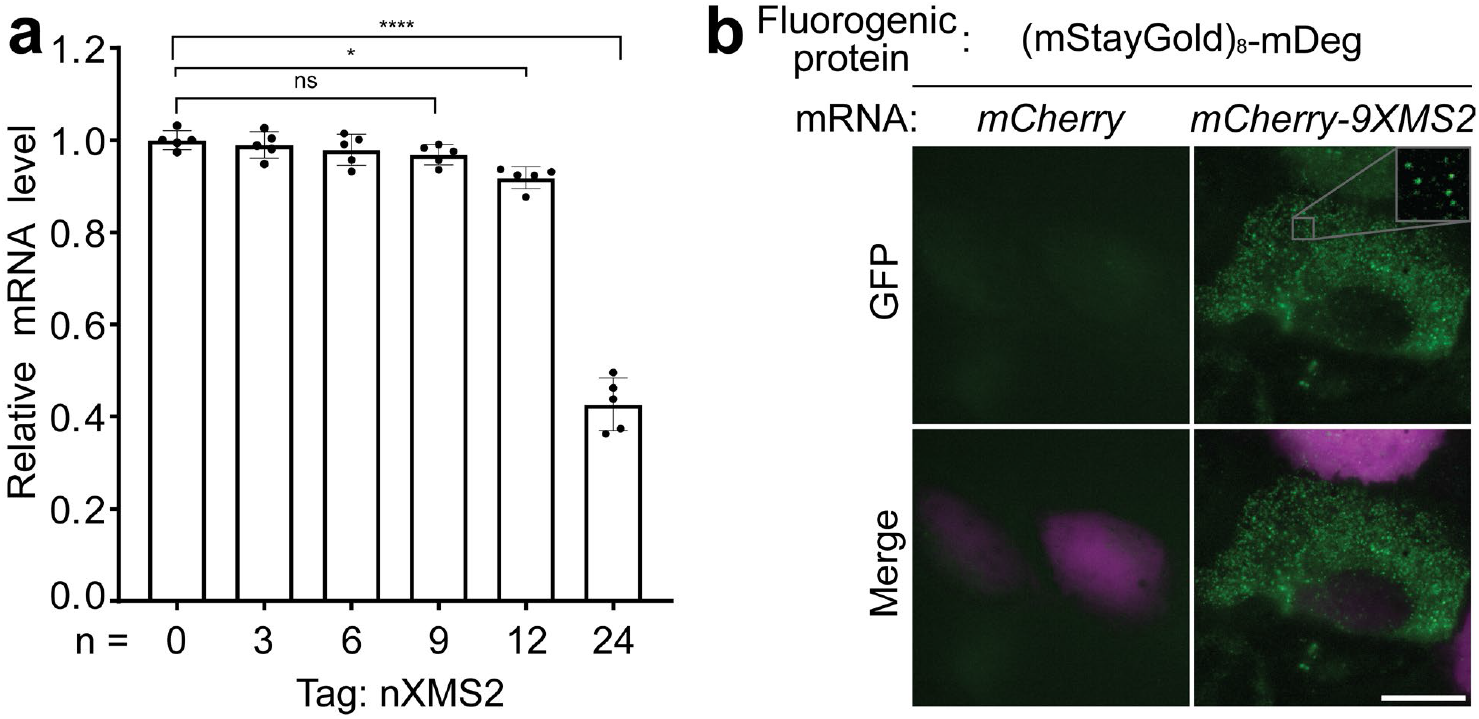
9XMS2, a short RNA imaging tag with minimal perturbation of RNA stability, for single mRNA imaging. **(a)** 9XMS2 has minimal perturbation on the tagged *mCherry* mRNA stability in living cells. To design a short MS2 tag with minimal perturbation to RNA stability, we performed RT-qPCR to quantify the levels of *mCherry* mRNA transcripts tagged with different repeat numbers (n = 3, 6, 9, 12, and 24, respectively) of MS2, and compared these to the same *mCherry* mRNA transcript without any tag (n = 0) in HEK293T cells. Our results showed that 9XMS2 has minimal perturbation on the tagged *mCherry* mRNA stability in living cells. (*n* = 5 independent cell cultures). **(b)** 9XMS2 enables single mRNA imaging when combined with (mStayGold)_8_-mDeg. To image mRNA tagged with 9XMS2, we coexpressed an *mCherry-9XMS2* mRNA reporter with (mStayGold)_8_-mDeg as the fluorogenic protein, in U2OS cells. We observed mobile green-fluorescent puncta in the cytosol of the U2OS cells as also shown in Supplementary Video 3. Importantly, these mobile green-fluorescent puncta were not observed in cells coexpressing (mStayGold)_8_-mDeg and an *mCherry* mRNA without any MS2 tag. Scale bar, 20 μm.

We then asked whether 9XMS2 can be used for single mRNA imaging in living cells. A key to imaging mRNA in living cells is to recruit enough fluorophores to the tagged mRNA while minimizing the background fluorescence from fluorophores that are not bound to the mRNA. To achieve this, we constructed a plasmid expressing mDeg fused to eight concatenated mStayGold, a highly photostable fluorescent protein with high molecular brightness^36^. We used this (mStayGold)_8_-mDeg to image an *mCherry* mRNA tagged with 9XMS2 in living U2OS cells. We observed mobile green-fluorescent puncta in the cytosol of the U2OS cells (Fig. 6b and Supplementary Video 3), likely reflecting single mRNAs. Importantly, these mobile green-fluorescent puncta were not observed in cells coexpressing (mStayGold)_8_-mDeg and an *mCherry* mRNA without any MS2 tag (Fig. 6b), suggesting that the green-fluorescent puncta are coming from the (mStayGold)_8_-mDeg bound to 9XMS2 in the tagged *mCherry* mRNA. Additionally, due to the high photostability of mStayGold, we were able to continuously track the dynamics of the 9XMS2 tagged *mCherry* mRNA for up to 40 seconds (Supplementary Video 3). Together, these results suggest that, when combined with (mStayGold)_8_-mDeg, a short MS2 tag (9XMS2) can be used for mRNA imaging with minimal perturbation of RNA stability.

## DISCUSSION

Overall, this study describes the development of two orthogonal RNA-regulated destabilization domains for live-cell three-color RNA imaging with minimal perturbation of RNA stability. These two RNA-regulated destabilization domains (termed mDeg and pDeg, respectively) were engineered from the most widely used RNA-RNA binding protein pairs, MS2-MCP and PP7-PCP, respectively. When fused to the C-terminus of fluorescent proteins, mDeg and pDeg can convert these constitutively on fluorescent proteins to RNA-regulated fluorogenic proteins. We showed that MS2-mDeg and PP7-pDeg can regulate the stability of a broad array of fluorescent proteins and HaloTag for RNA imaging. Due to their fluorogenicity and orthogonality, we showed that mDeg and pDeg can be combined with tDeg for simultaneous imaging the dynamics of three different mRNA species in living mammalian cells by transient transfection. Additionally, we identified 9XMS2 as a short tag for mRNA imaging without perturbing the tagged RNA’s stability.

RNA-induced fluorogenicity is an emerging concept that has been widely explored using fluorogenic RNA aptamers and small molecule dyes^29,30^. However, there have been limited examples of RNA-induced fluorogenicity in fluorescent proteins^13,37^. Here, we showed that RNA-induced fluorogenicity can be achieved by engineering stdMCP and stdPCP. Our design principle is similar to the Pepper-tDeg system, where the Arg-Arg-Arg-Gly degron is fused to the C-terminus of an RNA-binding protein such that RNA binding will block this degron from being recognized by the proteasomal machinery needed for proteolysis. Since the C-terminus of stdMCP and stdPCP is far away from their RNA-binding site, we applied circular permutation to generate a new C-terminus that is adjacent to their RNA-binding site (Supplementary Fig. 1). Our results suggest that the distance between the Arg-Arg-Arg-Gly degron and the RNA-binding site is critical for RNA-induced fluorogenicity (Fig. 1 and Supplementary Fig. 5). Future mutagenesis studies on optimizing the relative orientation between the RNA and the RNA-regulated destabilization domain may further improve its RNA-induced fluorogenicity.

During the development of pDeg, we unexpectedly found that all EYFP-fused circularly permuted stdPCP vairants showed yellow-fluorescent puncta, suggesting protein aggregation when expressed in cells (Supplementary Fig. 6). We found that this aggregation problem was not caused by binding to PP7 or circular permutation of the stdPCP protein. Additionally, we also observed similar yellow-fluorescent puncta when expressing the non-circularly permutated stdPCP fused to EYFP (EYFP-stdPCP) in cells. We hypothesized that the aggregation of stdPCP is due to the remaining residues in the FG loop. By removing these remaining FG loop residues (Val64 to Val78), we observed diffused yellow fluorescence without any observable aggregates, further confirming the remaining FG loop primarily contributes to the aggregation of stdPCP.

Unlikely conventional RNA hairpin methods based on MS2-MCP and PP7-PCP, MS2-mDeg and PP7-pDeg have low background fluorescence and thus does not require fusing an NLS to the fluorescent protein to decrease background fluorescence signals. Because of this, mDeg and pDeg can be readily used for imaging RNA with single-molecule resolution by transient transfection. Additionally, concatenated fluorescent proteins and self-labeling proteins with high molecular brightness and high photostability, such as mStayGold and HaloTag with the JF dyes, can be easily fused to mDeg and pDeg for live-cell RNA imaging. Importantly, these RNA-regulated fluorogenic proteins can be readily applied to imaging RNAs that are tagged with the widely used 24XMS2 tag and 24XPP7 tag.

Compared to the Pepper-tDeg system, MS2-mDeg can detect single mRNA in living cells more effectively. Using the same fluorogenic proteins, (mNeonGreen)_4_, we showed that MS2-mDeg can detect single mRNA in 94% of the transfected cells, which is higher than the 67% and 64% when using PP7-pDeg and Pepper-tDeg, respectively. This suggests a major improvement on the reliability for RNA detection using MS2-mDeg. On the other hand, the performance of PP7-pDeg is on par with the current system, Pepper-tDeg. Even with modest performance, PP7-pDeg can be combined with MS2-mDeg and Pepper-tDeg for two- and three-color RNA imaging in living mammalian cells.

Long RNA tags have been shown to destabilize the tagged RNAs in cells^14,15^. Having shorter RNA tags for imaging is critical to faithfully recapitulate the behavior of endogenous mRNA. With the fluorogenic property and the robustness of MS2-mDeg for RNA imaging, we showed that 9XMS2 can be used as a short tag with minimal perturbation of the tagged mRNA’s stability. Importantly, this is achieved through taking advantage of the fluorogenic property of MS2-mDeg by using a highly fluorescent and photostable fluorogenic protein, (mStayGold)_8_-mDeg.

Although these new RNA-regulated destabilization domains were used for live-cell RNA imaging in this study, the ability to use already widely adapted RNA aptamers, such as MS2 and PP7, to control protein stability would likely enable numerous applications in the field of RNA biology and synthetic biology. Conceivably, the MS2-mDeg and PP7-pDeg systems described here can be used for tethered function assays^38^ to elucidate the function of RNA-binding proteins without potential artifacts due to protein aggregations from overexpression of RNA-binding proteins with low complexity domains. Additionally, MS2-mDeg and PP7-pDeg can be readily combined with the CRISPR-Cas technology for gene expression regulation, genome imaging, and other synthetic biology applications.

## METHODS

### General methods and materials

All gene fragments used for constructing DNA plasmids used in this study were purchased from Twist Bioscience. For routine PCR amplifications, single-stranded synthetic DNA oligonucleotides were purchased from Integrated DNA Technologies, Phusion High-Fidelity PCR Master Mix (M0531L) was purchased from New England Biolabs. For restriction digest of PCR products and DNA plasmids, restriction endonucleases were purchased from New England Biolabs, and used according to the manufacturer’s recommended protocol. PCR products and digested DNA plasmids were purified using 0.8% TAE (Tris–acetate– ethylenediaminetetraacetic acid (EDTA) agarose gels and Thermo Scientific Gel Extraction kit (FERK0692). For DNA ligation reactions, Quick Ligation Kit (M2200L) was purchased from New England Biolabs. All DNA plasmids used in this study were propagated using chemically competent *Escherichia coli* (Agilent, 200315) then extracted using Thermo Scientific GeneJET Plasmid Miniprep Kit (FERK0503). Purified DNA plasmids were sent to Azenta Life Sciences or Plasmidsaurus for DNA sequencing to verify sequence identity.

### Cell culture and transfection

All mammalian cell lines used in this study, including HEK293T/17 (American Type Culture Collection (ATCC), CRL-11268), U2OS (ATCC, HTB-96), COS-7 (ATCC, CRL-1651), and HeLa (ATCC, CCL-2) cells, were cultured in Dulbeco’s modified Eagle’s medium (Thermo Fisher Scientific, 11995-065) supplemented with 10% fetal bovine serum (Corning 35-010-CV), 100 U ml^−1^ of penicillin and 100 μg ml^−1^ of streptomycin (Thermo Fisher Scientific, 15140122) under 37 °C with 5% CO_2_. For cell passage, TrypLE Express (Thermo Fisher Scientific, 12604013) was used for dissociating the adherent mammalian cells from culture flasks. All transfections in this study were carried out using FuGENE HD (Promega, E2312) according to the manufacturer’s recommended protocol. Cell culture media was changed to FluoroBrite™ DMEM (A1896701) prior to live-cell imaging.

### Live-cell fluorescence imaging of RNA in mammalian cells

DNA plasmids expressing all the mDeg- and pDeg-tagged fluorogenic proteins and HaloTag used in this study are based on a pcDNA3.1(+) vector backbone containing a miniCMV promoter followed by Kozak sequence (5’-GCCACC-3’). Gene fragments encoding all the fluorogenic proteins and HaloTag were purchased from Twist Bioscience. These genes fragments were cloned in the pcDNA3.1(+) vector using standard restriction enzyme cloning procedures.

DNA plasmids expressing different circular RNAs are based on a pAV U6+27 vector backbone containing the Tornado expression system, which was a gift from Samie Jaffrey (Addgene plasmid # 129405). Double-stranded synthetic DNA oligonucleotides encoding MS2 and PP7 were purchased from Integrated DNA Technologies and cloned into this circular RNA expression vector backbone containing an F30 folding scaffold^39^ using standard restriction enzyme cloning procedures.

DNA plasmids expressing reporter mRNA are based on a pcDNA3.1(+) vector backbone containing a CMV promoter followed by Kozak sequence (5’-GCCACC-3’). The (F30-2xPepper)_10_ tag was a gift from Samie Jaffrey (Addgene plasmid # 129403). The 24xMS2 tag was a gift from Robert Singer (Addgene plasmid # 84561). The 24xPP7 tag was a gift from Robert Singer (Addgene plasmid # 40652).

For fluorescence imaging of mDeg- and pDeg-tagged proteins with circular RNA, HEK293T (3.2 × 10^5^), U2OS (2.5 × 10^5^), COS-7 (2.5 × 10^5^), HeLa (2.5 × 10^5^) cells were seeded into 35 mm glass bottom dishes precoated with poly-D-lysine (Metek, P35GC-1.5-14-C), respectively, and cultured overnight. On the next day, cells were transfected using FuGENE HD according to the manufacturer’s recommended protocol. Specifically, 1400 ng of DNA plasmids encoding for fluorogenic protein fused to mDeg or pDeg were mixed with 1400 ng of DNA plasmids encoding for circular RNA for transfecting each 35 mm dish. The cells were imaged two days after transfection. Cell culture media was changed to FluoroBrite™ DMEM (A1896701) prior to live-cell imaging. For fluorogenic proteins containing HaloTag, a manufacturer’s recommended amount of the JF646 ligand was incubated with the transfected cells for at least one hour prior to imaging.

For fluorescence imaging of mRNAs, U2OS (2.5 × 10^5^) cells were seeded into 35 mm glass bottom dishes precoated with poly-D-lysine (Metek, P35GC-1.5-14-C) and culture overnight.

On the next day, cells were transfected using FuGENE HD according to the manufacturer’s recommended protocol. Specifically, a total of 2800 ng of DNA plasmids were used for transfecting each 35 mm dish. For cells that were co-transfected with more than one DNA plasmid, the amount of each plasmid equals to 2800 ng divided by the number of DNA plasmids used. The cells were imaged two days after transfection. Cell culture media was changed to FluoroBrite™ DMEM (A1896701) prior to live-cell imaging. For fluorogenic proteins containing HaloTag, a manufacturer’s recommended amount of the JF646 ligand was incubated with the transfected cells for at least one hour prior to imaging.

For live-cell fluorescence imaging, a Nikon Ti2-E epifluorescence inverted microscope equipped with a Prime BSI Express sCMOS monochrome camera, a Lumencor SOLA V-NIR Light Engine for Epifluorescence light source, and a temperature and CO_2_ stage top incubator (Tokai Hit) was used. The NIS-Elements Advanced Research software (Nikon) was used for controlling the microscope and camera. For live-cell imaging, a 20×/0.80-NA (numerical aperture) or a 60×/1.42-NA oil immersion objective (Nikon) with TYPE 37 immersion oil was used. A GFP filter cube (with excitation filter 470 ± 20 nm, dichroic mirror 495 nm (long pass), and emission filter 525 ± 25 nm) was used for detecting fluorogenic proteins with green fluorescence emissions. A YFP filter cube (with excitation filter 500 ± 10 nm, dichroic mirror 515 nm (long pass), and emission filter 535 ± 15 nm) was used for detecting fluorogenic proteins with yellow fluorescence emissions. A dsRed filter cube (with excitation filter 545 ± 15 nm, dichroic mirror 570 nm (long pass), and emission filter 620 ± 30 nm) was used for detecting fluorogenic proteins with red fluorescence emissions. A Cy5 filter cube (with excitation filter 620 ± 30 nm, dichroic mirror 660 nm (long pass), and emission filter 700 ± 37.5 nm) was used for detecting fluorogenic proteins with far-red fluorescence emissions. A DAPI filter cube (with 395 ± 12.5 nm excitation filter, 425 nm (long pass) dichroic mirror, and 460 ± 25 nm emission filter) was used for detecting the Hoechst-stained nuclei and mTagBFP2. ImageJ was used for analyzing fluorescence images. Cellular fluorescence intensity was calculated by measuring the mean fluorescence signal a cell’s area and subtracting background based on average signal of culture media. Normalized fluorescence was calculated by dividing the cell fluorescence intensity of each cell to the averaged cell fluorescence of the whole cell population.

### Quantitative PCR with reverse transcription

Total cellular RNAs were extracted with a GeneJET RNA Purification Kit (Thermo Fisher Scientific, K0731) according to the manufacturer’s instructions. The purified RNAs were then reverse-transcribed to the cDNAs using SuperScript™ IV First-Strand Synthesis kit (Invitrogen, 18080051) with an oligo-dT and random hexamer primers according to the manufacturer’s instructions. Quantitative PCR was performed to measure the relative mRNA level by mixing the cDNA with SYBR Green Supermix (Thermo Fisher Scientific, A25741). The amplification conditions were as follows: 50°C for 2 mins, 95°C for 2 min, 40 cycles of 95°C for 15 s, 56°C for 15 s and 72°C for 1 min. The mCherry and GAPDH primers used for the amplification were listed in Supplementary Table 1. *mCherry* expression measured were normalized to that of the housekeeping genes GAPDH using 2^-(Cq mCherry - Cq GAPDH)^, where Cq is the amplification cycle measured by qPCR. Biological replicates were tested.

## Statistical analysis

All data are expressed as means ± s.d. with sample sizes (n) listed for each experiment. Statistical analyses were performed using Excel (Microsoft) and Prism (Graphpad). One-way ANOVA and unpaired, two-tailed, Student’s t-test were used to analyze significant differences between the group means.

## Supplementary Information

is linked to the online version of the paper.

## Acknowledgements

This work was supported by the UMass startup funds (T.G.P., O.A., and J.W.). We thank Dr. James Chambers for technical support on imaging experiments. We also thank other members of the Wu lab for useful comments and suggestions.

## Author Contributions

J.W. conceived the project. J.W., T.G.P., and O.A. designed experiments. T.G.P. and O.A. performed experiments. J.W., T.G.P., and O.A. analyzed data and wrote the manuscript.

## Competing Financial Interests

The authors declare no competing financial interests.

## Notes

### Competing Interest Statement

The authors have declared no competing interest.

