## Supplementary Information for "Orthogonal RNA-regulated destabilization domains for three-color RNA imaging with minimal RNA perturbation"

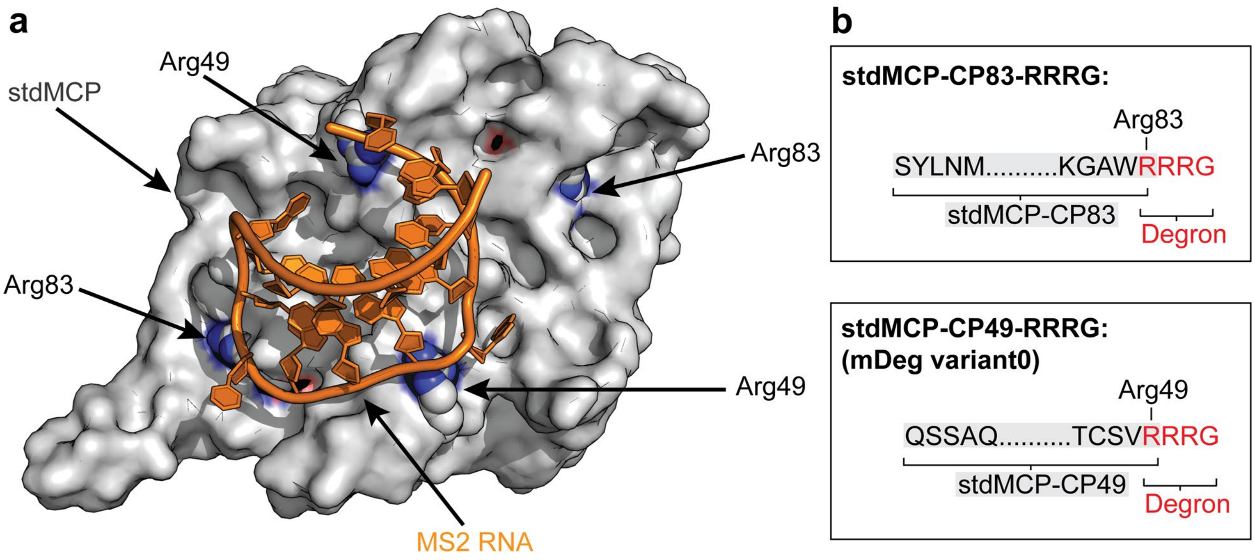


**Supplementary Figure 1. Initial designs of an MS2-regulated destabilization domain.**

Shown is the crystal structure of MS2-stdMCP (PDB: 2BU1). (**a**) and our initial designs for converting stdMCP to an MS2-regulated destabilization domain (**b**). (**a**) The MS2 RNA is depicted in orange and the stdMCP protein is depicted in grey with side chains of Arg49 and Arg83 highlighted in each MCP monomer. To engineer an MS2-regulated destabilization domain, we circularly permuted stdMCP at Arg49 and Arg83, respectively, and inserted “Arg-Arg-Gly” at the new C-terminus to generate the full “Arg-Arg-Arg-Gly” degron. We chose Arg49 and Arg83 for circular permutation because they are close to the MS2 binding site, and thus, binding of the MS2 RNA may block the Arg-Arg-Arg-Gly degron from recruitment of the proteasomal machinery needed for proteolysis. (**b**) Partial sequence of our designs for an MS2-regulated destabilization domain. Amino acid sequence of circularly permutated stdMCP is highlighted in grey, the “Arg-Arg-Arg-Gly” degron sequence is in red.

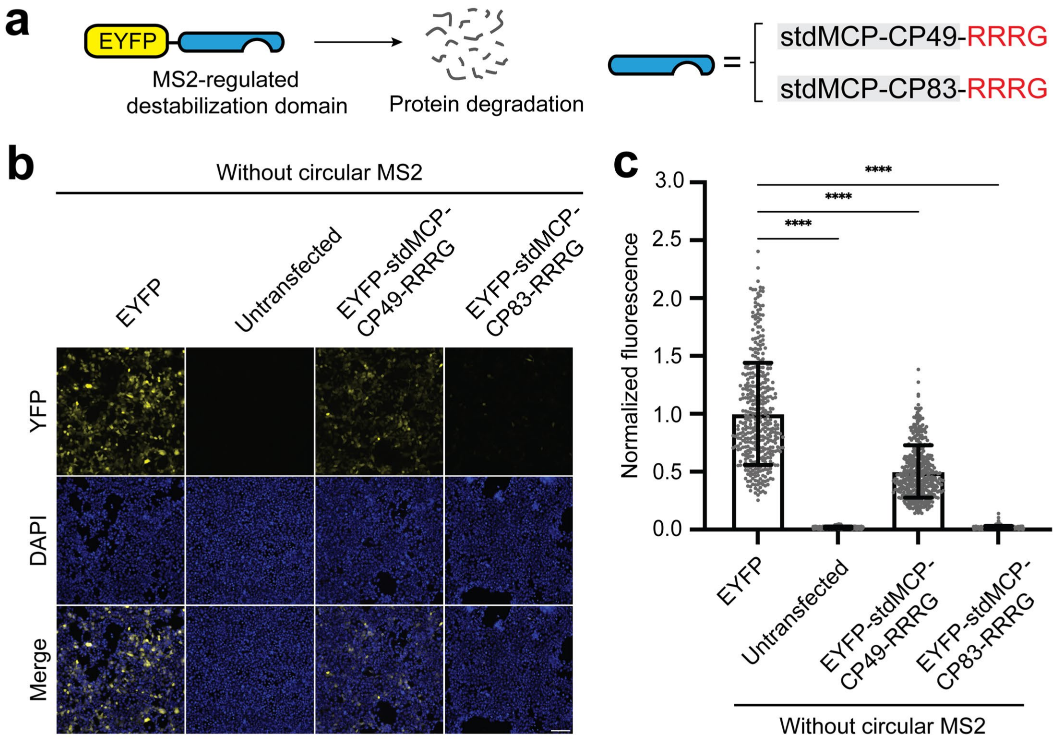


**Supplementary Figure 2. Compared to stdMCP-CP49-RRRG, stdMCP-CP83-RRRG reduces the stability of EYFP to a much greater extent.**

In Supplementary Figure 1, we showed our initial designs for engineering an MS2-regulated destabilization domain, termed stdMCP-CP49-RRRG, and stdMCP-CP83-RRRG, respectively. Here, we tested whether fusing stdMCP-CP49-RRRG and stdMCP-CP83-RRRG, respectively, to a protein confers protein instability. (**a**) A schematic showing the proteins tested in these experiments. We fused stdMCP-CP49-RRRG and stdMCP-CP83-RRRG to the C-terminus of enhanced yellow fluorescent protein (EYFP), respectively, and expressed this fusion protein in HEK293T cells. (**b**, **c**) When expressed in HEK293T cells, EYFP-stdMCP-CP49-RRRG and stdMCP-CP83-RRRG showed yellow fluorescence that is 50% and 2% of cells expressing EYFP, respectively. All cells were stained with Hoechst dye. Scale bar, 100 µm. In (**c**), normalized average cellular yellow fluorescence of individual cells is plotted. ($n = 3$ independent cell cultures). Values are means ± s.d.

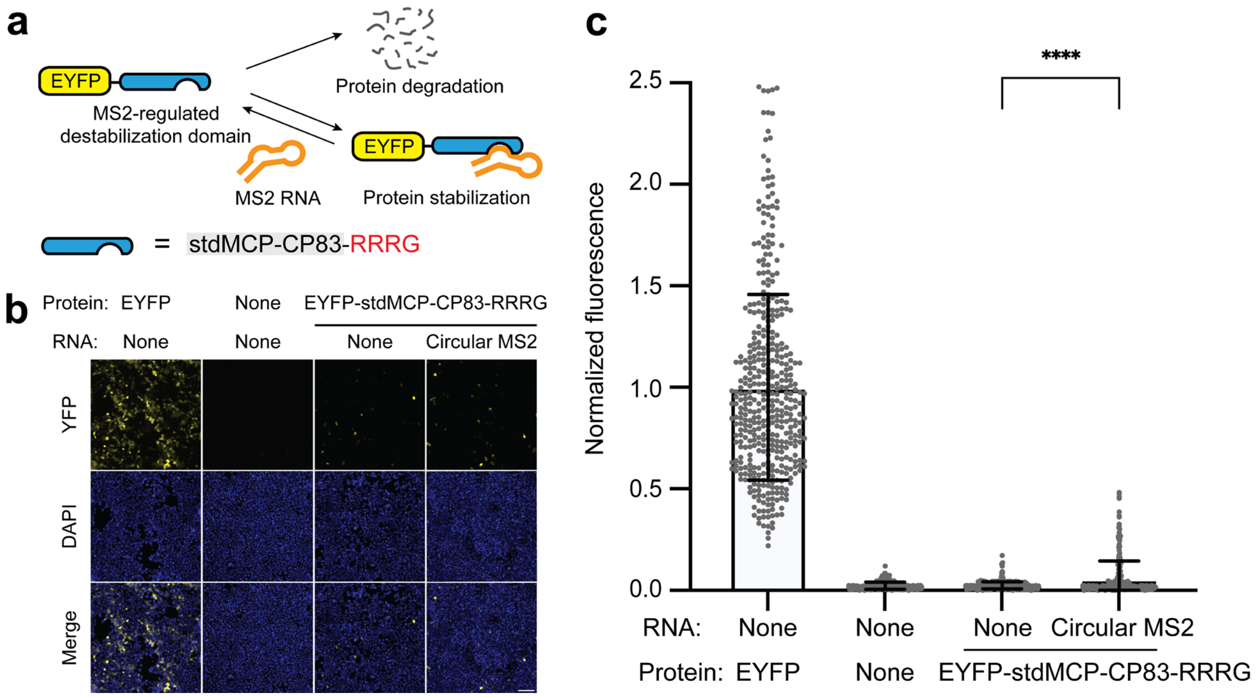


**Supplementary Figure 3. MS2 cannot regulate the stability of stdMCP-CP83-RRRG.**

In Supplementary Figure 2, our results suggest that stdMCP-CP83-RRRG confers instability to EYFP most efficiently. Here, we tested whether the destabilization effect from stdMCP-CP83-RRRG can be inhibited by binding to MS2. (**a**) A schematic showing the protein and RNA tested in these experiments. (**b**, **c**) We coexpressed EYFP-stdMCP-CP83-RRRG with the MS2 RNA and a control RNA in HEK293T cells. We observed that coexprression of the circular MS2 RNA only led to a very limited increase in yellow fluorescence compared to the circular control RNA. This suggests that MS2 cannot regulate the stability of stdMCP-CP83-RRRG. All cells were stained with Hoechst dye. Scale bar, 100 μm. Normalized average cellular yellow fluorescence of individual cells is plotted. ($n$ = 3 independent cell cultures). Values are means ± s.d.

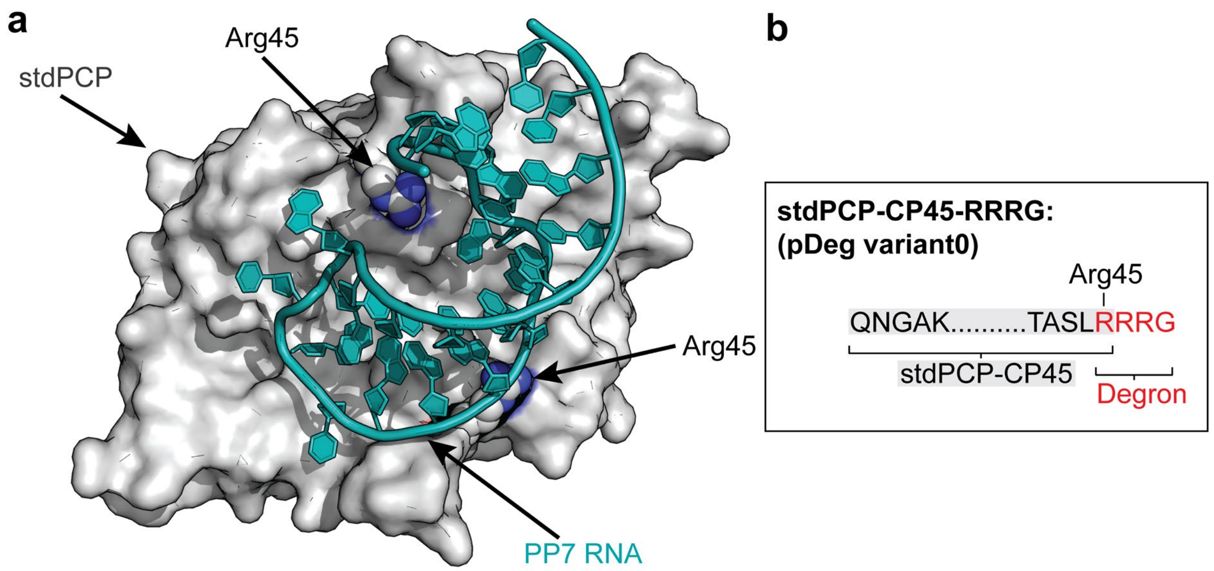


**Supplementary Figure 4. Initial design of a PP7-regulated destabilization domain.**

Shown is the crystal structure of PP7-stdPCP (PDB: 2QUX). (**a**), and our initial design for converting stdPCP to a PP7-regulated destabilization domain (**b**). (**a**) The PP7 RNA is depicted in tiffany green and the stdPCP protein is depicted in grey with side chains of Arg45 (equivalent to Arg49 in stdMCP) highlighted in each PCP monomer. Similar to the initial design of mDeg, we circularly permuted stdPCP at Arg45 and inserted “Arg-Arg-Gly” at the new C-terminus to generate the full “Arg-Arg-Arg-Gly” degron. We termed this stdPCP-CP45-RRRG, also known as pDeg variant0 in Supplementary Fig. 5. The idea is that the binding of the PP7 RNA to stdPCP-CP45-RRRG may block the Arg-Arg-Arg-Gly degron from recruitment of the proteasomal machinery needed for proteolysis. (**b**) Partial sequence of our design for a PP7-regulated destabilization domain. Amino acid sequence of circularly permutated stdPCP is highlighted in grey, the “Arg-Arg-Arg-Gly” degron sequence is in red.

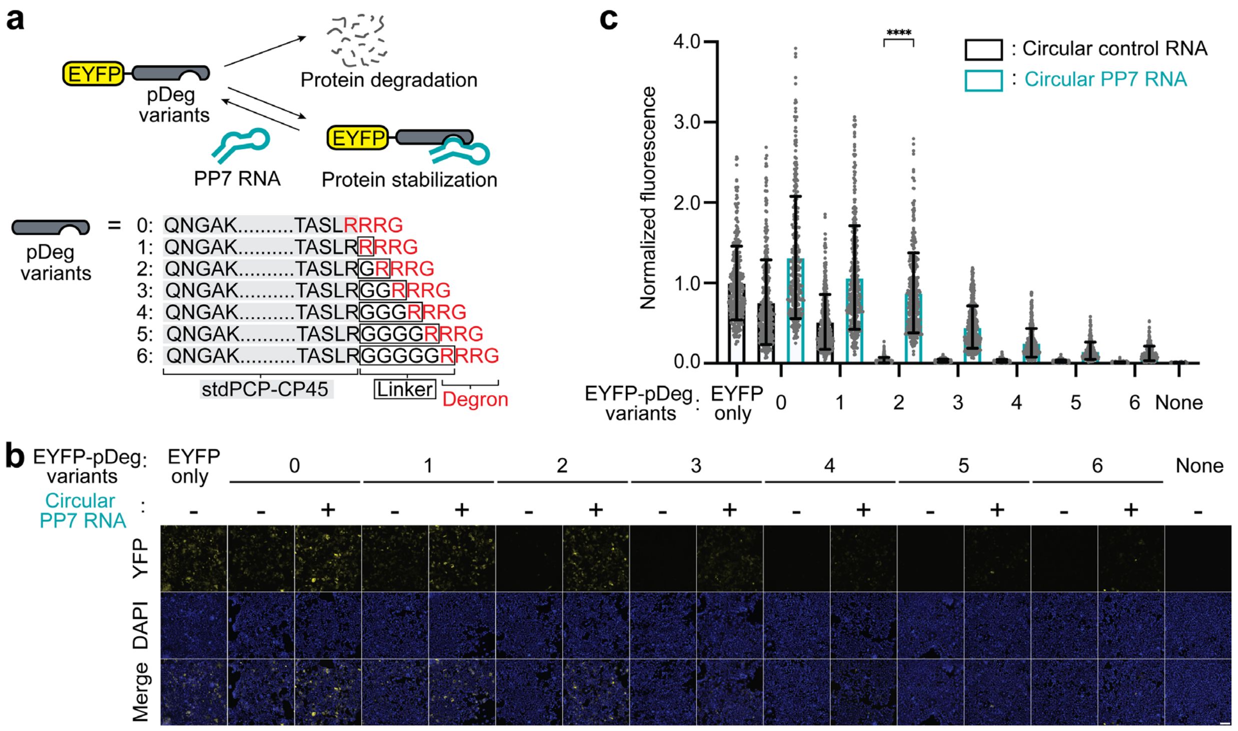


**Supplementary Figure 5. The stability of pDeg variants can be regulated by the PP7 RNA.**

To construct a PP7-regulated destabilization domain, we used a similar strategy used for developing mDeg. Specifically, we titrated linker length from one to six amino acids between stdPCP-CP45 and the Arg-Arg-Arg-Gly degron, resulting in seven pDeg variant (0-6) as shown in (**a**). To test whether PP7 can regulate the stability of these different pDeg variants (0-6), we coexpressed each EYFP-pDeg variant with and without circular PP7 in HEK293T cells, respectively. We found that stdPCP-CP45-RRRG (pDeg variant0) showed limited degradation efficiency as it exhibited 76% of yellow fluorescence compared to EYFP. The addition of linkers between stdPCP-CP45 and the Arg-Arg-Arg-Gly degron led to decreases of yellow fluorescence in the absence of circular PP7. When PP7 was coexpressed, we found that pDeg variant2 containing an “Arg-Gly” linker showed the highest fold change in yellow fluorescence of 22-fold, as shown in (**b**) and (**c**). All cells were stained with Hoechst’s dye. Scale bar, 100 µm. Normalized average cellular yellow fluorescence of individual cells is plotted in (**c**). ($n = 3$ independent cell cultures). Values are means ± s.d.

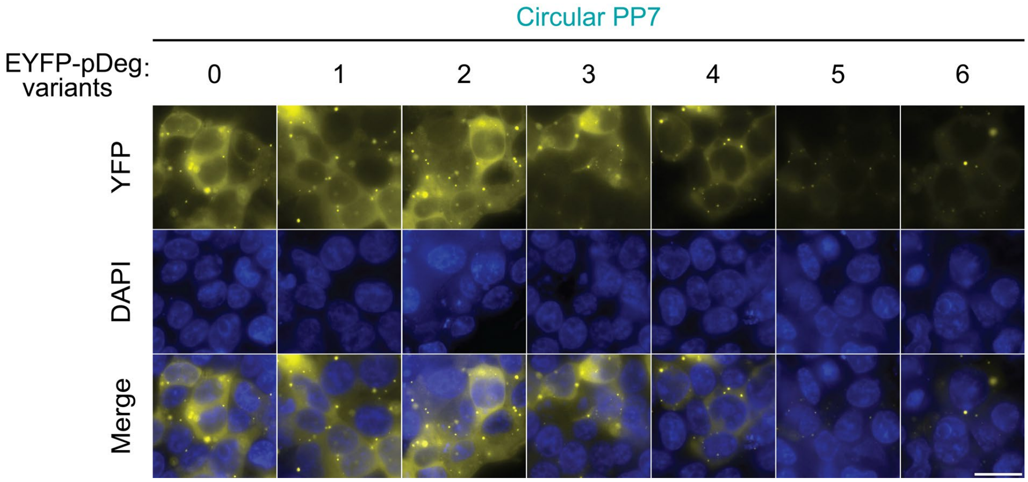


**Supplementary Figure 6. All EYFP-pDeg variants showed yellow-fluorescent puncta when coexpressed with circular PP7 in cells.**

In Supplementary Figure 5, we showed that the stability of most pDeg variants can be regulated by the PP7 RNA. However, we unexpectedly observed yellow-fluorescent puncta in all EYFP-pDeg variants when coexpressed with circular PP7 in HEK293T cells. The yellow-fluorescent puncta suggest aggregation of these EYFP-pDeg variants when in complex with PP7. All cells were stained with Hoechst's dye. Scale bar, 20 μm.

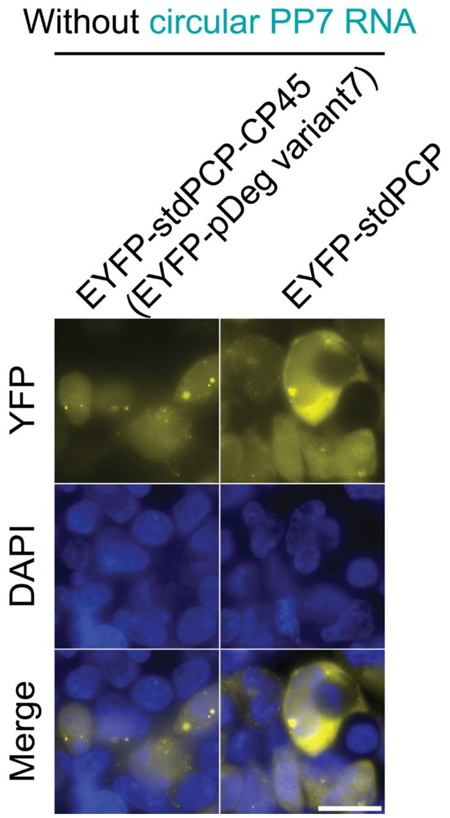


**Supplementary Figure 7. The aggregation of the EYFP-pDeg variant2-PP7 complex does not come from the PP7 RNA or circular permutation of pDeg variant2.**

In Supplementary Figures 5 and 6, we showed that PP7 can regulate the stability of most pDeg variants in cells. In particular, EYFP-pDeg variant2 showed the highest fluorescence fold change in response to binding to circular PP7 in HEK293T cells. However, all the EYFP-pDeg variants also showed aggregation in cells.

Here, we sought to eliminate this aggregation phenomenon by first asking whether the yellow-fluorescent puncta of EYFP-pDeg variant2 is due to aggregation of the circular PP7 RNA. To test this, we generated a pDeg variant2 lacking the Arg-Arg-Arg-Gly degron (termed pDeg variant7). The idea is that pDeg variant7 should be stable in cells without any circular PP7 RNA, due to the lack of the Arg-Arg-Arg-Gly degron. If the yellow-fluorescent puncta observed in Supplementary Figures 6 are due to binding to the circular PP7 RNA, we should not observe any of these puncta in cells expressing EYFP-pDeg variant7. However, we found that EYFP-pDeg variant7 showed similar yellow-fluorescent puncta as in cells expressing the EYFP-pDeg variant2-PP7 complex. Therefore, this suggests that the aggregation of the EYFP-pDeg variant2-PP7 complex does not come from the circular PP7 RNA.

Since circular permutation could change the energetics of transient partial unfolding in proteins, which may lead to protein aggregation, we asked whether the aggregation of the EYFP-pDeg variant2-PP7 complex is due to aggregation of pDeg variant2. To test this, we expressed the non-circularly permuted stdPCP fused to EYFP in HEK293T cells. Unexpectedly, we observed that EYFP-stdPCP itself can cause aggregations in cells. This suggests that the yellow-fluorescent puncta are not caused by circular permutation of stdPCP. All cells were stained with Hoechst's dye. Scale bar, 20 μm.

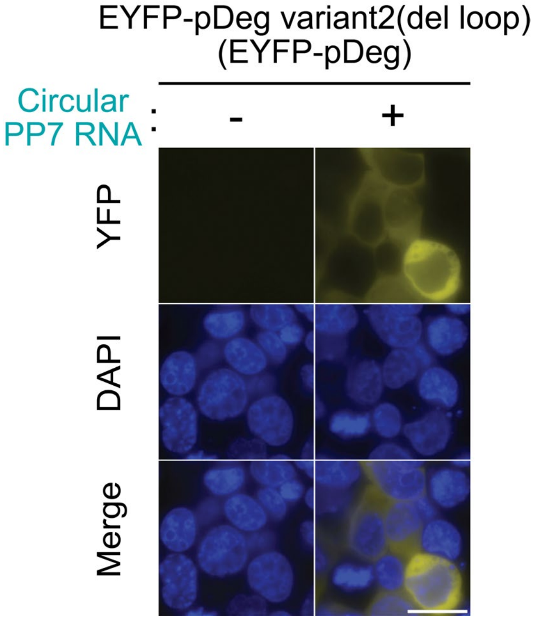


**Supplementary Figure 8. Removal of the remaining amino acids (Cys67 to Phe74) in the FG loop of PCP eliminates aggregation of the EYFP-pDeg variant2-PP7 complex in cells.**

In Supplementary 7, we showed that the aggregation of the EYFP-pDeg variant2-PP7 complex does not come from the PP7 RNA or circular permutation of pDeg variant2. Here, we asked whether the remaining FG loop residues (Cys67 to Phe74) in each PCP monomer could be the cause of aggregation. To test this, we replaced the remaining FG loop (Val64 to Val78) with a single Gly residue in pDeg variant2 and termed this pDeg variant2(del loop). When coexpressed with circular PP7, EYFP-variant2(del loop) did not show any yellow-fluorescent puncta in HEK293T cells. These results suggest that the fluorescent puncta of EYFP-pDeg variant2 was caused by the remaining residues in the FG loop. We termed pDeg variant2 (del loop) as "pDeg." All cells were stained with Hoechst's dye. Scale bar, 20 μm.

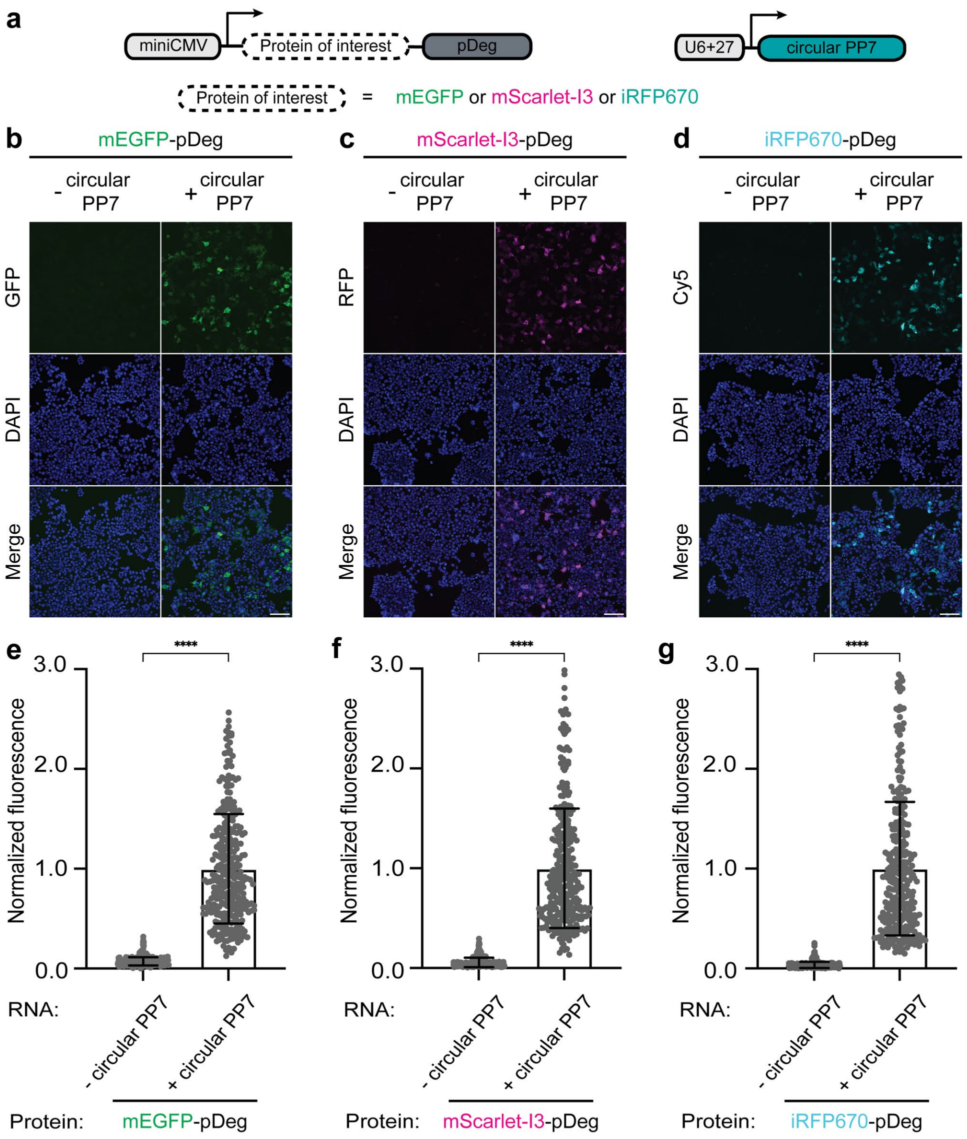


**Supplementary Figure 9. PP7-pDeg confers PP7-dependent regulation to various fluorescent proteins.**

(**a**) Scheme of plasmids that encode protein of interest fused to pDeg, and circular PP7 RNA, respectively. Protein of interest includes mEGFP, mScarlet-I3, and iRFP670.

(**b-d**) PP7-pDeg can regulate the stability of various fluorescent proteins. To test whether the PP7 RNA stabilizes different proteins fused to pDeg, we imaged HEK293T cells expressing mEGFP (**b**), mScarlet-I3 (**c**), and iRFP670 (**d**) fused to pDeg with and without circular PP7, respectively. In each case, there was a considerable increase in fluorescence. All cells were stained with Hoechst's dye. Scale bar, 100 µm.

(**e-g**) Summary data of normalized fluorescence of pDeg-fused fluorescent proteins with and without circular PP7 as in (**b-d**). Normalized average cellular fluorescence of individual cells is plotted. ($n = 3$ independent cell cultures). Values are means ± s.d.

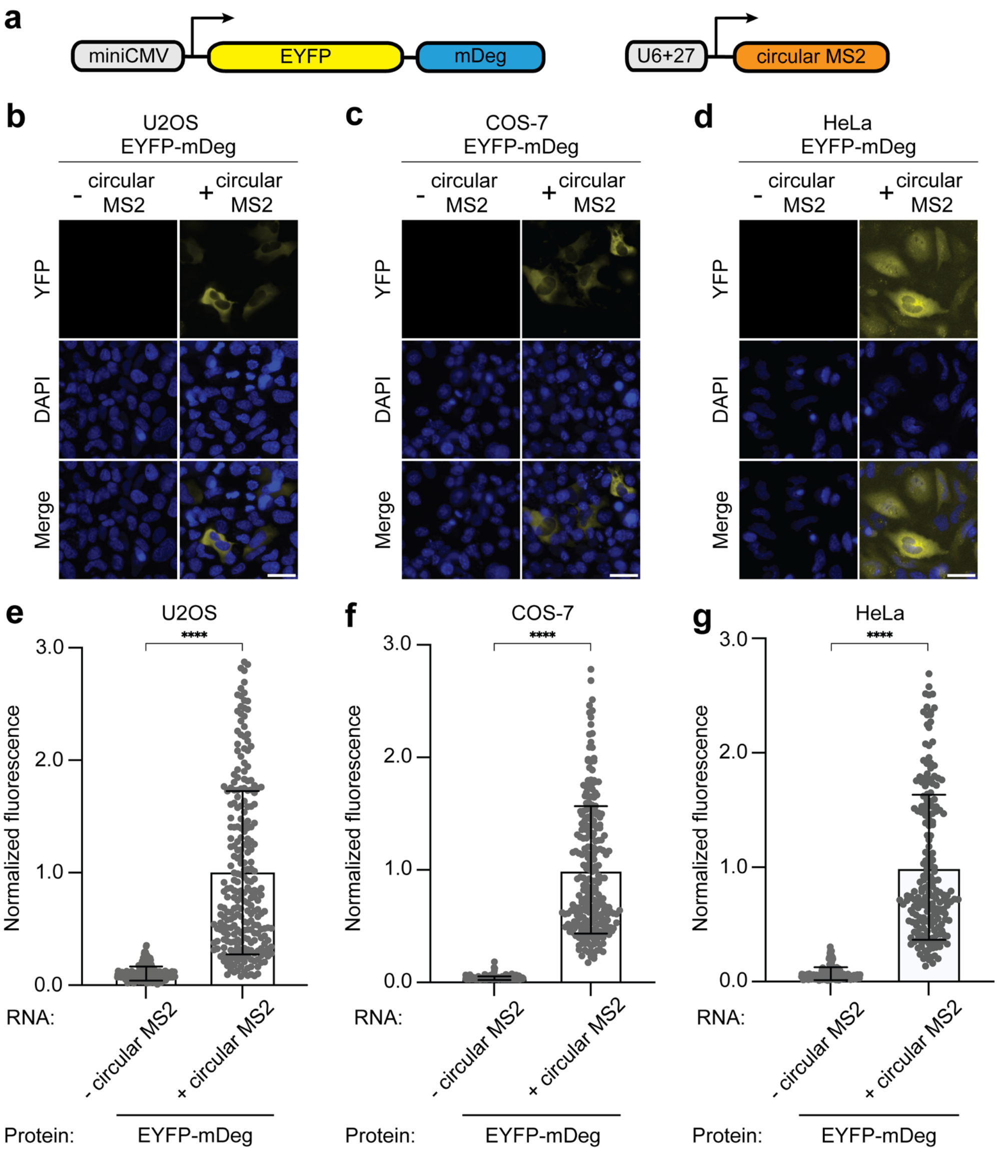


**Supplementary Figure 10. mDeg can be regulated by the MS2 RNA in diverse mammalian cell types.**

In Fig. 1, we showed that EYFP-mDeg can be regulated by MS2 in HEK293T cells. Here, we tested whether mDeg can be regulated by MS2 in various mammalian cell lines. To test this, we expressed EYFP-mDeg in U2OS (**b**), COS-7 (**c**), and HeLa cells (**d**), respectively. In each case, the EYFP-mDeg was coexpressed with and without circular MS2. Similar to what we observed in HEK293T cells, EYFP-mDeg showed minimal fluorescence in all three cell lines without MS2. However, the yellow fluorescence intensity of EYFP-mDeg was considerably increased only when circular MS2 was coexpressed. Thus, these results suggest that mDeg can be regulated by MS2 in various mammalian cell lines. All cells were stained with Hoechst's dye. Scale bar, 40 µm.

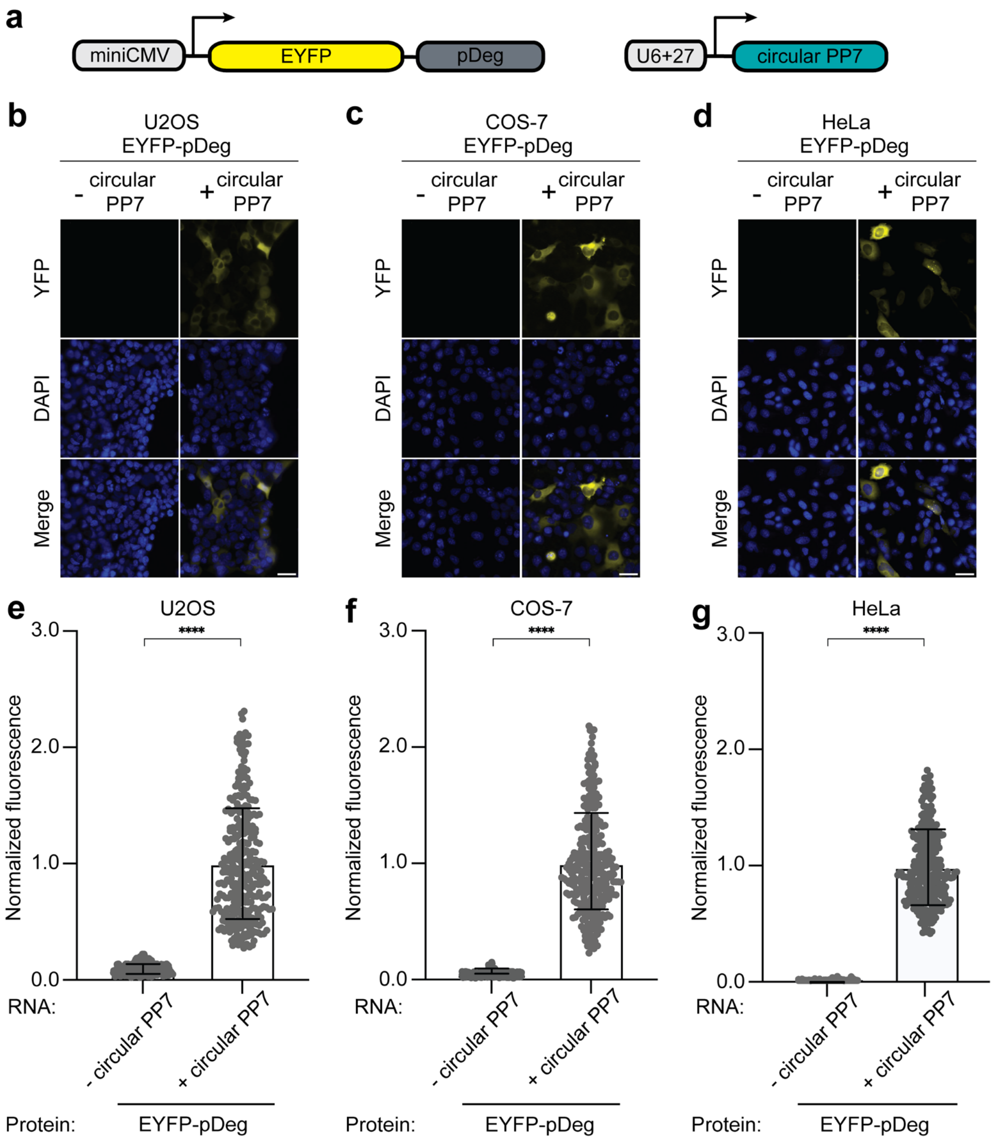


**Supplementary Figure 11. pDeg can be regulated by the PP7 RNA in diverse mammalian cell types.**

In Fig. 2, we showed that EYFP-pDeg can be regulated by PP7 in HEK293T cells. Here, we tested whether pDeg can be regulated by PP7 in various mammalian cell lines. To test this, we expressed EYFP-pDeg in U2OS (**b**), COS-7 (**c**), and HeLa cells (**d**), respectively. In each case, the EYFP-pDeg was coexpressed with and without circular PP7. Similar to what we observed in HEK293T cells, EYFP-pDeg showed minimal fluorescence in all three cell lines without PP7. However, the yellow fluorescence intensity of EYFP-pDeg was considerably increased only when circular PP7 was coexpressed. Thus, these results suggest that pDeg can be regulated by PP7 in various mammalian cell lines. All cells were stained with Hoechst's dye. Scale bar, 40 μm.

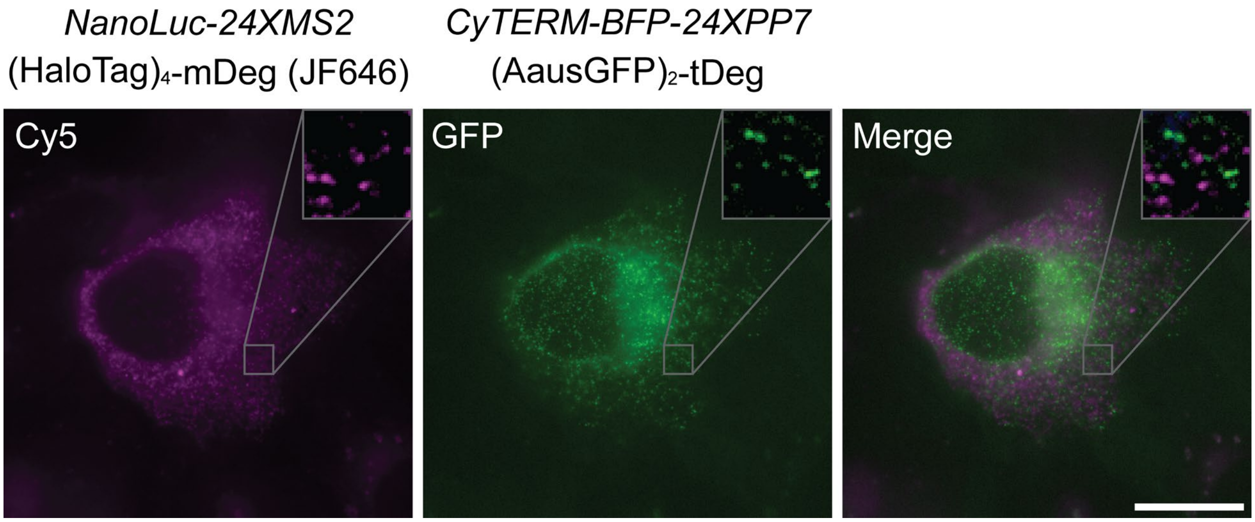


**Supplementary Figure 12. MS2-mDeg and PP7-pDeg enable two-color RNA imaging in living mammalian cells.**

In Fig. 5, we showed MS2-mDeg, PP7-pDeg, and Pepper-tDeg are fully orthogonal to each other in mammalian cells. Based on this, we sought to use these systems to image three different RNA species in living cells at the same time. As a first step, we asked whether we could use MS2-mDeg and PP7-pDeg for two-color RNA imaging. To test this, we expressed two reporter plasmids expressing *Nanoluc* tagged with 24XMS2 and *CyTERM-mTagBFP2* tagged with 24xPP7, respectively. In the same cells, we coexpressed $(HaloTag)_4$-mDeg (with the JF646 ligand) and $(AausGFP)_2$-pDeg as the fluorogenic proteins. We observed distinct but non-overlapping fluorescent puncta in both the GFP channel and Cy5 channel, reflecting individual mRNAs. This suggests that MS2-mDeg and PP7-pDeg can be used for tracking the dynamics of two RNA species in the same cell at the same time. Scale bar, 20 µm.

**Supplementary Table 1. ssDNA oligo primers used in RT-qPCR**

| | |
|---|---|
| mCherry fw | CACGAGTTCGAGATCGAGGG |
| mCherry rv | CAAGTAGTCGGGGATGTCGG |
| GAPDH fw | AAGGTCGGAGTCAACGGATT |
| GAPDH rv | TGAGGTCAATGAAGGGGTCA |

**Supplementary Video 1. MS2-mDeg, PP7-pDeg, and Pepper-tDeg enable three-color RNA imaging in living cells**
To simultaneously image three different RNA species in U2OS cells, we coexpressed three mRNA reporters: *Nanoluc-24XMS2*, *CDK6-24XPP7*, and *CyTERM-mTagBFP2-(F30-2xPepper)*$_{10}$ with three fluorogenic proteins: (HaloTag)$_4$-mDeg (with the JF646 ligand) (cyan), tdTomato-pDeg (magenta), and (mNeonGreen)$_4$-tDeg (green). We observed mobile fluorescent puncta in the cytosol reflecting cellular mRNAs. Scale bar, 20 μm.

**Supplementary Video 2. MS2-mDeg can be combined with photostable fluorogenic proteins for continuous RNA imaging for an extended period.**
To test whether mDeg can be combined with photostable fluorophores for continuous RNA imaging for an extended period of time. We coexpressed (HaloTag)$_4$-mDeg (with the JF646 ligand) and a *NanoLuc-24XMS2* mRNA in U2OS cells. We observed mobile far-red fluorescent puncta with continuous image acquisition for 120 seconds with limited photobleaching. Scale bar, 20 μm.

**Supplementary Video 3. 9XMS2, a short RNA imaging tag with minimal perturbation of RNA stability, for single mRNA imaging.**
To image mRNA tagged with 9XMS2, we coexpressed an *mCherry-9XMS2* mRNA reporter with (mStayGold)$_8$-mDeg as the fluorogenic protein, in U2OS cells. We observed mobile green-fluorescent puncta in the cytosol of the U2OS cells, reflecting single mRNA molecules. Scale bar, 20 μm.
